# Site-specific phosphorylation of Huntingtin exon 1 recombinant proteins enabled by the discovery of novel kinases

**DOI:** 10.1101/2020.07.23.217968

**Authors:** Anass Chiki, Jonathan Ricci, Ramanath Hegde, Luciano A. Abriata, Andreas Reif, Hilal A. Lashuel

## Abstract

Posttranslational modifications (PTMs) within the first 17 amino acids (Nt17) of exon1 of the Huntingtin protein (Httex1) play important roles in modulating its cellular properties and functions in health and disease. In particular, phosphorylation of threonine and serine residues (T3, S13, and/or S16) has been shown to inhibit Htt aggregation *in vitro* and inclusion formation in cellular and animal models of Huntington’s disease (HD). In this manuscript, we describe a new and simple methodology for producing milligram quantities of highly pure wild type or mutant Httex1 proteins that are site-specifically phosphorylated at T3 or at both S13 and S16. This advance was enabled by 1) the discovery and validation of novel kinases that efficiently phosphorylate Httex1 at S13 and S16 (TBK1), at T3 (GCK) or T3 and S13 (TNIK and HGK); and, 2) the development of an efficient methodology for producing recombinant native Httex1 proteins using a SUMO-fusion expression and purification strategy. As proof of concept, we demonstrate how this method can be applied to produce Httex1 proteins that are both site-specifically phosphorylated and fluorescently labeled or isotopically labeled. Together, these advances should increase access to these valuable tools and expand the range of methods and experimental approaches that can be used to elucidate the mechanisms by which phosphorylation influences Httex1 structure, aggregation, interactome and function(s) in health and disease.

## Introduction

Huntington’s disease (HD) is a devastating neurodegenerative disease for which there are currently no effective treatments or disease-modifying therapies. HD is characterized by severe motor symptoms such as chorea, bradykinesia, loss of motor control, rigidity [1-3], and difficulties with speech and swallowing [4]. As the disease progresses, HD patients experience cognitive impairments, personality change, and depression [5-7]. HD is a monogenic disease that is caused by a CAG expansion in the first exon (exon 1) of the *HTT* gene [8, 9], which is translated into a polyglutamine (polyQ) repeat in the huntingtin protein HTT [10]. Individuals with polyQ repeat lengths that extend beyond the pathogenic threshold of ≥36 glutamine residues go on to develop HD. The higher the number of polyQ repeats, the more severe the symptoms are, and the earlier is the age of disease onset [11]. Although the exact mechanisms underpinning neurodegeneration in HD remain unclear, converging evidence suggests that polyQ expansions increase the propensity of HTT proteins to aggregate and form pathological inclusions in a polyQ-dependent manner [12]; the longer the polyQ repeats, the higher is the propensity of HTT to aggregate and form inclusions [13, 14]. The formation of HTT aggregates and inclusions has been linked to cellular dysfunction and degeneration via different cellular mechanisms [15-17].

Increasing evidence suggests that nuclear HTT inclusions in *postmortem* tissues of HD patients [18] are formed as a result of the misfolding and aggregation of N-terminal HTT fragments of varying lengths, rather than the full-length HTT protein [19]. One of the most studied fragments corresponds to exon 1 of HTT (Httex1), which contains the polyQ domain and is generated by aberrant splicing [20] and possibly proteolytic processing of the protein [21]. Overexpression of mutant Httex1 containing polyQ repeats ranging from 80 to 175 glutamine residues is sufficient to induce a robust HD-like phenotype and pathology in various animal models (mice, Drosophila) [17, 22, 23], as well as in cell culture models of HD [17, 24]. *In vitro*, mutant Httex1 exhibits polyQ-dependent aggregation and forms amyloid-like fibrils in a concentration-dependent manner [25, 26].

Posttranslational modifications (PTMs) in the first 17 N-terminal amino acids (Nt17) of Httex1 can dramatically affect HTT aggregation, subcellular targeting, clearance, and toxicity [14, 27, 28]. Phosphorylation, acetylation [27, 29], ubiquitination and SUMOylation [28] have been shown to occur in the Nt17 domain, with indications that some of these modifications coexist on the same molecule [29]. Given the reversible nature of these modifications, we hypothesized that they could act as a molecular switch for regulating many aspects of HTT, including its structure, interactome, and cellular properties. Therefore, a better understanding of the mechanisms by which these PTMs influence HTT structure, aggregation, and cellular properties may offer new avenues for the development of more effective disease-modifying strategies. Indeed, several lines of evidence suggest that Nt17 PTMs could reverse the deleterious effects caused by polyQ expansions. For example, mutating both residues S13 and S16 in the Nt17 region to aspartate to mimic phosphorylation was shown to inhibit Httex1 aggregation, sufficiently modify the aggregation properties of full-length HTT and protect against mutant HTT-induced toxicity in a transgenic model of HD [30]. Additionally, mutant HTT was shown to be hyperphosphorylated at S13 and S16 in *STHdh* cells and HD mice [31, 32]. Furthermore, restoring Nt17 phosphorylation induces K9 acetylation and promotes HTT clearance by the proteasome and lysosome [29]. Finally, we recently showed that the levels of T3 phosphorylation (pT3) inversely correlated with polyQ repeat length in both preclinical models of HD and samples from HD patients [27, 33].

Until recently, studies on the effect of Nt17 PTMs have relied on the use of mutations to mimic PTMs. These include the substitution of serine and threonine residues by the acidic residues aspartic acid or glutamic acid to mimic phosphorylation [27, 30] or replacing lysine with glutamine to mimic lysine acetylation [34]. These approaches have several limitations that have precluded a more accurate understanding of the role of PTMs in regulating HTT biology and its role in HD, including the fact that PTM mimetics do not fully capture the size, charge state, or dynamic nature of a *bona fide* PTMs. For example, phosphomimetics allow only partial mimicking of phosphorylation and do not reproduce the dynamic nature of this modification or its effects on the structural properties of Nt17^[25^. It is, therefore, not surprising to observe conflicting findings concerning the effects of phosphomimetic on HTT aggregation when using different animal models [27, 30, 35].

To address these limitations, we recently developed a semisynthetic methodology that enables the site-specific introduction of single or multiple PTMs in WT and mutant. Using this approach, we gained new insight into how Httex1 aggregation and conformation are affected by Nt17 phosphorylation, acetylation, or the cross-talk between these two types of PTMs [33, 36-39]. Moreover, the ability to generate site-specifically modified Httex1 proteins enabled us to develop very sensitive assays to detect and quantify phosphorylated Httex1 in complex samples for biomarker discovery [33, 40]. However, the current semisynthetic methods for the production of phosphorylated Httex1 proteins have some limitations: 1) they are time-consuming and require advanced technical capabilities in protein chemical synthesis; 2) introducing PTMs beyond residue 9 requires the introduction of a non-native Gln-18 to Ala mutation to enable native chemical ligation; and 3) they are not suitable for the production of both modified and isotopically labeled Httex1 proteins for structural studies using NMR and other methods.

To overcome these challenges, it is crucial to first identify the enzymes responsible for Nt17 phosphorylation and then develop efficient *in vitro* phosphorylation conditions that allow site-specific phosphorylation at the desired residues. Towards this goal, we performed kinase screening using a library of 298 kinases and identified several kinases that phosphorylate HTT efficiently and specifically at T3 or both S13 and S16. Next, we took advantage of the specificity and efficiency of these kinases to develop efficient methods that allowed the production of milligram quantities of homogenously phosphorylated recombinant Httex1 at T3 or both S13 and S16. This was enabled by the recently developed SUMO-based Httex1 expression and purification strategy [41], enabling the generation of highly pure milligram quantities of Httex1 proteins. To demonstrate the versatility of these methods, we present examples that illustrate how they could be used to produce fluorescently and isotopically labeled phosphorylated untagged WT (23Q) and mutant (43Q) Httex1 proteins. Furthermore, using NMR, we provide some preliminary results on the cross-talk between S13 and S16 and how phosphorylation could influence the structural properties of Httex1. Together, these advances should pave the way for future studies to elucidate the effects of phosphorylation on the interactome, structural, and cellular properties of Httex1 that were previously not possible.

## Results and Discussion

### Identification and validation of kinases that phosphorylated Nt17 at T3, S13, and S16

To enable the site-specific enzymatic phosphorylation of Httex1 at T3, S13 or S16, we first needed to identify enzymes that phosphorylate HTT at these sites. Towards this goal, we performed a screening using the *In vitro* Kinase and Phosphopeptide Testing (IKPT) services from Kinexus [42], where Httex1-23Q and the Nt17 peptide were used as substrates to test a panel of 298 purified serine-threonine kinases. This led to the identification of several enzymes that phosphorylate Nt17 and Httex1 at both S13 and S16 (TBK1) or at T3 (GCK, TNIK, and HGK), respectively. We recently described the validation of TBK1 and demonstrated that it phosphorylates S13 and S16 efficiently and specifically *in vitro* and S13 in cells and *in vivo [43]*. Herein, we present for the first time the discovery and validation of the kinases that phosphorylate T3.

The top kinase hits that phosphorylated T3 in the context of Httex1 included MAP4K2 (GCK), MAP4K4 (HGK), and TNIK (Figure 1). These kinases are part of the STE20 kinase family and are involved in cellular signal transduction, including apoptosis, the cell cycle, and cell growth [44]. To determine the efficiency and specificity of these kinases, we performed *in vitro* phosphorylation reactions in which these kinases were co-incubated with Httex1-23Q (Figure 1) and monitored the extent of phosphorylation and number of phosphorylation sites by Electrospray Ionisation Mass Spectrometry (ESI/MS) and western blot (WB) using our well-tested and validated phospho-antibodies [37, 40, 45] specific for pT3, pS13, or pS16. As shown in Figure 1, GCK phosphorylated Httex1-23Q mainly at one site, as indicated by the +80 Da mass shift observed by ESI/MS (Figure 1-A). In contrast, HGK and TNIK phosphorylated Httex1-23Q at multiple sites, as illustrated by the appearance of two new peaks in the ESI/MS spectra that correspond to the phosphorylation of one or two sites (Figure 1-A). Western blot analysis using pT3-, pS13- and pS16-specific antibodies enabled us to determine which sites were phosphorylated. As shown in Figure 1-B, GCK phosphorylates mainly at T3, as indicated by a strong positive WB signal with the pT3 antibody (Figure 1-B), and the absence of any bands in the WB developed using pS13 and pS16 antibodies. On the other hand, both T3 and S13 phosphorylation were detected with their respective antibodies when Httex1- 23Q was coincubated with HGK or TNIK (Figure 1-B), which was consistent with the ESI/MS results showing phosphorylation at multiple sites in Nt17. Interestingly, the coexpression of each of these kinases with Httex1- 16Q-eGFP in HEK 293 cells did not result in significant phosphorylation (Figure S1). Nevertheless, these findings did not exclude the potential of using the newly discovered kinases as tools to prepare phosphorylated Httex1 *in vitro*. The efficiency and specificity of GCK in phosphorylating T3 *in vitro* suggest that it could be used as a valuable tool for the preparation of Httex1 proteins that are site-specifically and homogeneously phosphorylated at T3 (pT3).

**Figure 1.**
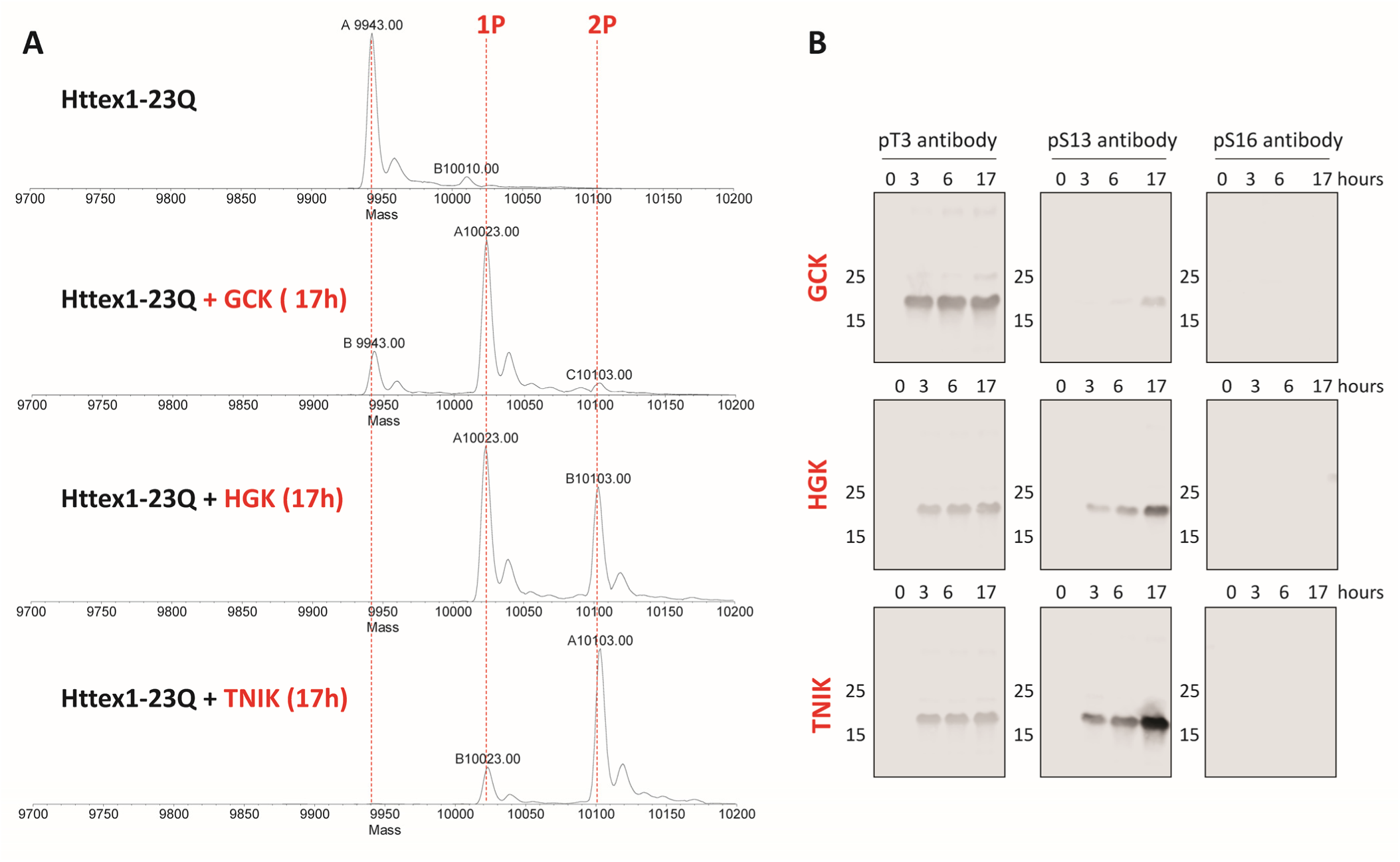
Validation of Httex1 phosphorylation by GCK, HGK, and TNIK. **(A)** ESI/MS analysis of Httex1-23Q before and after the phosphorylation reaction with GCK, HGK, or TNIK. **(B)** Western blot analysis of Httex1-23Q phosphorylation by GCK, HGK, and TNIK monitored over-time by homemade antibodies specific to pT3, pS13, or pS16.

Having shown that both GCK and TBK1 phosphorylate wild-type Httex1 with 23Q, we next sought to determine if they could phosphorylate mutant Httex1 (Httex1-43Q) with the same efficiency and specificity. Due to its high aggregation propensity, Httex1-43Q was tested at a lower concentration (20 M). As shown in Figure 2, MALDI analysis indicated that GCK partially phosphorylated (∼30%) mutant Httex1 at T3 (Figure 2-A); even after a prolonged incubation time (16 hours), the extent of phosphorylation did not significantly increase (Figure 2-A). However, TBK1 rapidly phosphorylated Httex1-43Q at both S13 and S16 (Figure 2-B). After 4 hours, most of the protein exhibited phosphorylation at two sites, and only a small portion of the protein was singly phosphorylated (Figure 2-B). After 16 hours of incubation, the level of monophosphorylated Httex1-43Q decreased (Figure 2-B). However, as shown in Figure 2-C, we observed significant Httex1-43Q aggregation for both kinases, as indicated by the appearance of higher molecular weight species in the WB (highlighted in red in Figure 2-C). These species were shown to form after 2 hours of the phosphorylation reaction (Figure 2-C). The formation of aggregates was caused by the conditions of the kinase reaction, which required a high protein concentration as well as incubation of the reaction mixture at 30°C. Thus, the high propensity of mutant HTT to aggregate *in vitro* precluded the generation of site-specific phosphorylated mHttex1 proteins through direct *in vitro* phosphorylation of the native Httex1 proteins by GCK and TBK1.

**Figure 2.**
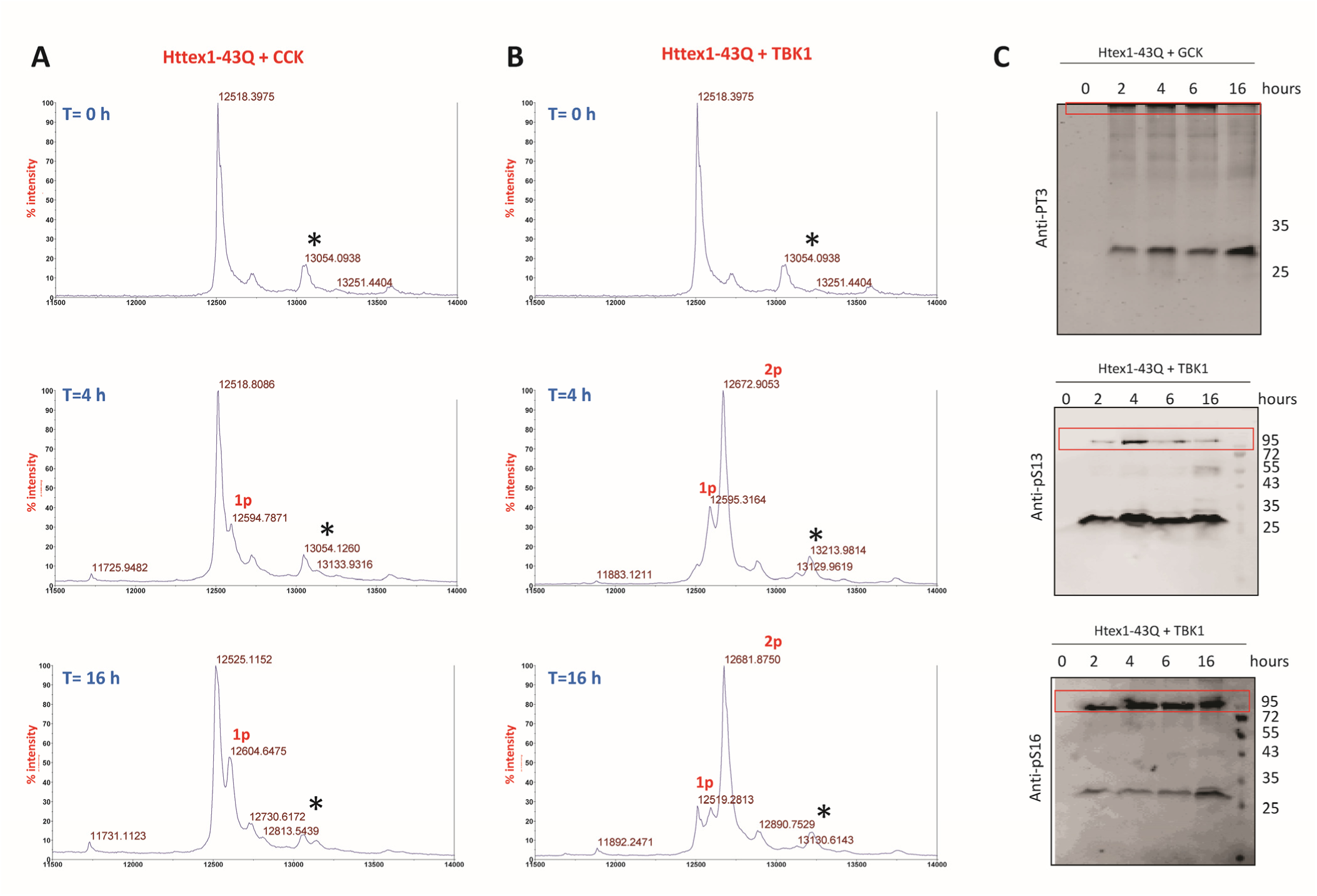
*In vitro* phosphorylation of Httex1-43Q by GCK or TBK1. Representative MALDI spectra of Httex1-43Q over-time phosphorylation by GCK **(A)** or by TBK1 **(B)** (T= 0 h is the same for both GCK and TBK1, sinapinic acid adducts are highlighted with a star). **(C)** Representative western blot of Httex1-43Q phosphorylation over time by GCK and TBK1 using pT3-, pS13-, or pS16-specific antibodies. Formed aggregates were observed during the phosphorylation reaction (highlighted by a red box).

### SUMO-based strategy for the generation of phosphorylated WT and mutant Httex1

To overcome the problems posed by the poor solubility of mutant Httex1, we took advantage of recent advances made by our group that enabled the efficient production of milligram quantities of highly pure WT and mutant Httex1 using the SUMO fusion strategy [41]. We showed that the fusion of SUMO to mutant Httex1 improves the expression of these proteins, increases their solubility, and facilitates their handling and purification. Therefore, we explored the possibility of performing the *in vitro* phosphorylation reaction directly on the mutant SUMO- Httex1 fusion protein (Scheme 1) with the idea that the SUMO tag can later be cleaved from Httex1 by the enzyme ULP1 rapidly to yield site-specifically phosphorylated native Httex1 proteins (Scheme 1). Therefore, we expressed Httex1 with 23 or 43 glutamine residues (Httex1-23Q or Httex1-43Q, respectively) containing a His6-SUMO tag fused to the N-terminus (SUMO-Httex1-23Q and SUMO-Httex1-43Q), as previously described [41]. After cell lysis, the fusion proteins were subjected to immobilized metal affinity chromatography (IMAC) (Figure S2-A and S2-B). Both SUMO-Httex1-23Q and SUMO-Httex1-43Q (SUMO-Httex1-Qn, n=23/43) were isolated by nickel affinity chromatography (Figure S2-A and S2-B, respectively), and the elution was analyzed by SDS-PAGE, as shown in Figure-S2-A and S2-B. Then, the fractions containing SUMO-Httex1-Qn were pooled and analyzed by ESI/MS and Ultra Performance Liquid Chromatography (UPLC) (Figure 3-A and 3-B). Although SDS-PAGE showed the presence of some impurities in the sample, ESI/MS showed the presence of mainly the SUMO-Httex1-Qn proteins (Figure- 3-A and 3-B). This could be because these protein impurities that do not ionize well on ESI/MS. Such impurities are usually removed during RP-HPLC purification. The UPLC analysis showed a broad single peak corresponding to the fusion proteins, as was previously shown [41]. To investigate the efficiency of the *in vitro* phosphorylation of SUMO-Httex1-Qn, SUMO-Httex1-23Q and SUMO-Httex1-43Q were both incubated with TBK1 and GCK, and the reaction was monitored by ESI/MS. Both kinases phosphorylated the WT and mutant fusion proteins, as evidenced by the appearance of additional phospho groups (+80 Da) by ESI/MS (Figure S3-A). After 6 hours, GCK induced complete phosphorylation of both the WT and mutant Httex1 at a single site (Figure S3-A). In the case of TBK1, a mixture of 2 and 3 phosphorylation sites was observed (Figure S3-A). Bearing in mind that TBK1 phosphorylates only two sites in Nt17 (S13 and S16) [43], we hypothesized that there are other potential TBK1 phosphorylation sites within the sequence of the SUMO tag. To reveal the actual state of phosphorylation, we performed a cleavage reaction of the SUMO tag by ULP1 on an analytical scale and analyzed the cleaved phosphorylated Httex1. Indeed, when analyzing the cleavage product from the reaction of SUMO-Httex1-23Q with GCK (6 hours of incubation), we observed that only 50% of Httex1-23Q was phosphorylated (Figure S3-B). On the other hand, the cleavage of SUMO-Httex1-23Q phosphorylated by TBK1 (6 hours of incubation) showed 100% phosphorylation at S13 and S16 only (Figure S3-B), confirming our hypothesis that the third phosphorylation occurs on the SUMO protein. Consequently, we established that it is essential to perform analytical SUMO cleavage to monitor phosphorylation for all our future applications. Moreover, to achieve complete phosphorylation of the desired sites, the *in vitro* phosphorylation reactions were extended to 17 hours, especially for GCK kinase. To verify that the phosphorylation conditions and extended incubation time did not affect the stability of SUMO-Httex1-43Q, we followed the phosphorylation by GCK or TBK1 using UPLC (Figure S2-C). For both kinases, the conditions did not affect the intensity of the SUMO-Httex1-43Q peak even after overnight incubation at 30°C (Figure S2-C), suggesting the absence of aggregation under these conditions.

**Figure 3.**
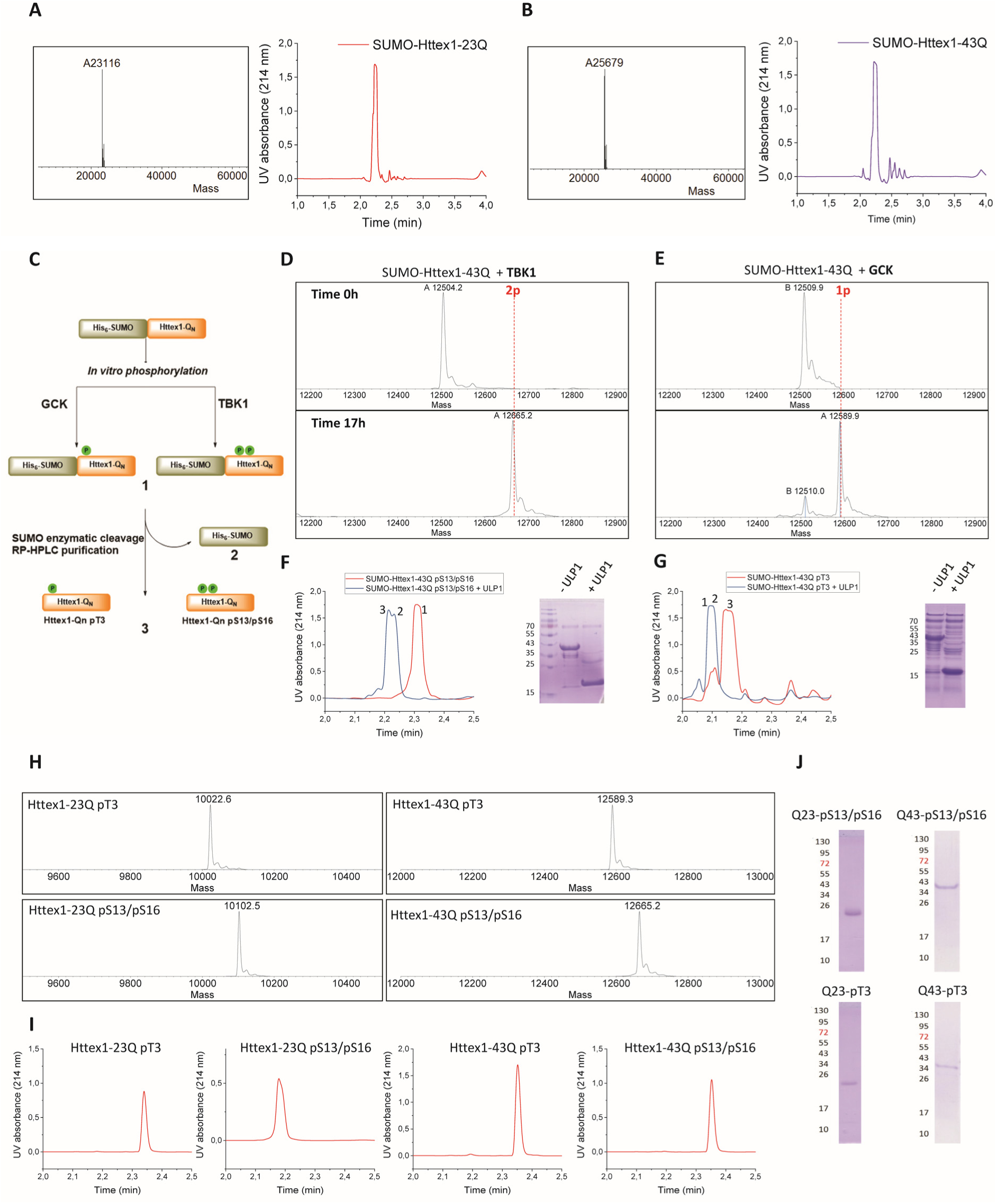
SUMO-based strategy for the *in vitro* quantitative production of phosphorylated Httex1 23Q and 43Q using TBK1. **(A-B)** Characterization of SUMO- Httex1-23Q **(A)** and SUMO-Httex1-43Q **(B)** by ESI/MS (left) and UPLC (right). **(C)** Schematic overview of the method used for phosphorylation of SUMO-Httex1- Qn. Monitoring of the phosphorylation reaction of SUMO-Httex1-43Q with TBK1 **(D)** and GCK **(E)** by ESI/MS after an analytical SUMO cleavage. Monitoring of the cleavage of the SUMO tag by ULP1 from phosphorylation reaction of SUMO-Httex1-43Q with TBK1 **(F)** or GCK **(G)**. Purity characterization of Httex1-23Q pT3, Httex1-43Q pT3, Httex1-23Q pS13/pS16, and Httex1-43Q pS13/pS16 by **(H)** ESI/MS, **(I)** UPLC and **(J)** SDS-PAGE. (Expected molecular weights for the proteins are listed in **Table S1**).

Having optimized and validated the phosphorylation of SUMO-Httex1-Qn, we performed a 10 mg preparative phosphorylation of SUMO-Httex1-23Q and SUMO-Httex1-43Q with GCK and TBK1. After overnight incubation, TBK1 phosphorylated SUMO-Httex1-23Q (Figure S4-A) and SUMO-Httex1-43Q (Figure 3-D) completely at both S13 and S16, as indicated by the addition of 2 phospho groups in the cleaved Httex1-Qn pS13/pS16 proteins (Figure S4-A and Figure 3-D). Once phosphorylation was confirmed, the SUMO tag was cleaved using ULP1, and the generation of the native phosphorylated Httex1-Qn proteins was monitored by UPLC and SDS-PAGE (Figure 3-F and Figure S4-C). After 15 min, cleavage of the SUMO tag was complete as indicated by the disappearance of the UPLC peak corresponding to SUMO-Httex1-Qn pS13/pS16 and the appearance of 2 new peaks for Httex1-Qn pS13/pS16 and the SUMO tag (Figure 3-F and Figure S4-C). This was further confirmed by SDS-PAGE, which showed the complete disappearance of SUMO-Httex1-Qn pS13/pS16 after the addition of ULP1 for 15 min (Figure 3-F and Figure S4-C). It should be noted that UPLC is a faster and more efficient tool to monitor this rapid cleavage reaction than SDS-PAGE. Next, the Httex1-Qn pS13/pS16 protein was separated from the SUMO tag and other impurities by reversed-phase HPLC (RP-HPLC) (Figure S4-E and S4-G). Httex1-Qn pS13/pS16 eluted first (∼22 min) and was separable from the SUMO tag, which eluted later (∼30 min) (Figure S4-E and S4-G). The fractions containing Httex1-Qn pS13/pS16 were pooled together, and the purities of Httex1-23Q pS13/pS16 and Httex1- 43Q pS13/pS16 were analyzed by ESI/MS, UPLC, and SDS-PAGE (Figure 3 H-J). Similarly, we used GCK to phosphorylate SUMO-Httex1-23Q and SUMO-Httex1-43Q at T3. Analysis of the overnight phosphorylation reaction by ESI/MS showed that the reaction was not complete (∼80% completion for the WT and mutant Httex1), as indicated by the detection of a mixture of nonphosphorylated and singly phosphorylated species, by ESI/MS, Figure 3-E and Figure S4-B. However, due to the appearance of a tiny population of doubly phosphorylated species (Figure S4-B), we decided not to push the phosphorylation reaction further with increased incubation time or the addition of more kinase in order to ensure single phosphorylation at T3. SUMO-Httex1-Qn phosphorylated by GCK was then incubated with ULP1 to remove the SUMO tag. After SUMO cleavage was confirmed by UPLC (Figure 3-G and Figure S4-D), the reaction mixture was subjected to RP-HPLC purification to separate Httex1-Qn pT3 from SUMO (Figure S4-F and S4-H). We were able to separate Httex1-pT3 23Q and Httex1-43Q pT3 from the nonphosphorylated and doubly phosphorylated species (highlighted in grey and green, respectively, in Figure S4- F and S4-H). Finally, the fractions containing our protein of interest were pooled together and analyzed by ESI/MS, UPLC, and SDS-PAGE, as indicated in Figure 3 H-J. This new protocol enabled us to enzymatically produce highly pure WT and mutant Httex1 phosphorylated at T3 or pS13/pS16 in milligram quantities for the first time. The complete characterization of the produced proteins (Httex1-23Q pT3, Httex1-23Q pS13/pS16, Httex1-43Q pT3, and Httex1-43Q pS13/pS16) by ESI/MS, UPLC, and Coomassie SDS PAGE is shown in Figure 3 H-J. With this optimized protocol in hand, we then moved to assess its utility to generate fluorescently and isotopically labeled and site-specifically phosphorylated Httex1 proteins for future use as valuable tools in structural studies (solution and solid-state NMR [46, 47]) and cellular studies to investigate the seeding and cell-to-cell transmission of HTT aggregates.

### Generation of fluorescently labeled (ATTO-565 maleimide) phosphorylated WT and mutant Httex1

Fluorescently labeled HTT proteins are valuable tools to understand the HTT structure and aggregation mechanisms *in vitro* [48] and to monitor its subcellular localization [49], dynamics of aggregation [50] and clearance, cell-to-cell transmission [51] and cellular properties in HD cellular models. GFP family fusion proteins are commonly used in HD cellular studies to track the formation of cytoplasmic and nuclear HTT aggregates and inclusions [35, 52-55]. However, these proteins are large compared to Httex1, which might significantly impact its structure and aggregation properties. Small molecule fluorescent probes present many advantages compared to fusion-labeled proteins; they are highly sensitive and stable and are likely to show less interference with the normal functions and interactome of the target protein. Amine-reactive probes can be used to label lysine residues; however, their labeling might result in the heterogenous preparation of labeled proteins; additionally, lysines within the Nt17 domain play an essential role in the amphipathic helix, and by consequence, any small modification might perturb the Nt17 structure. Furthermore, until now, semisynthesis was required to introduce fluorescent molecules in Httex1 with PTMs [39]. To overcome these challenges, we introduced a single-site mutation of proline to cysteine (P90C) at the C-terminus of SUMO-Httex1-Qn to enable labeling with Atto-565- maleimide (Figure 5-A). Maleimide fluorophores are highly specific for the thiol group of cysteine (P90C), and we showed previously that the reaction is rapid with the thiol of P90C [39].

**Figure 4.**
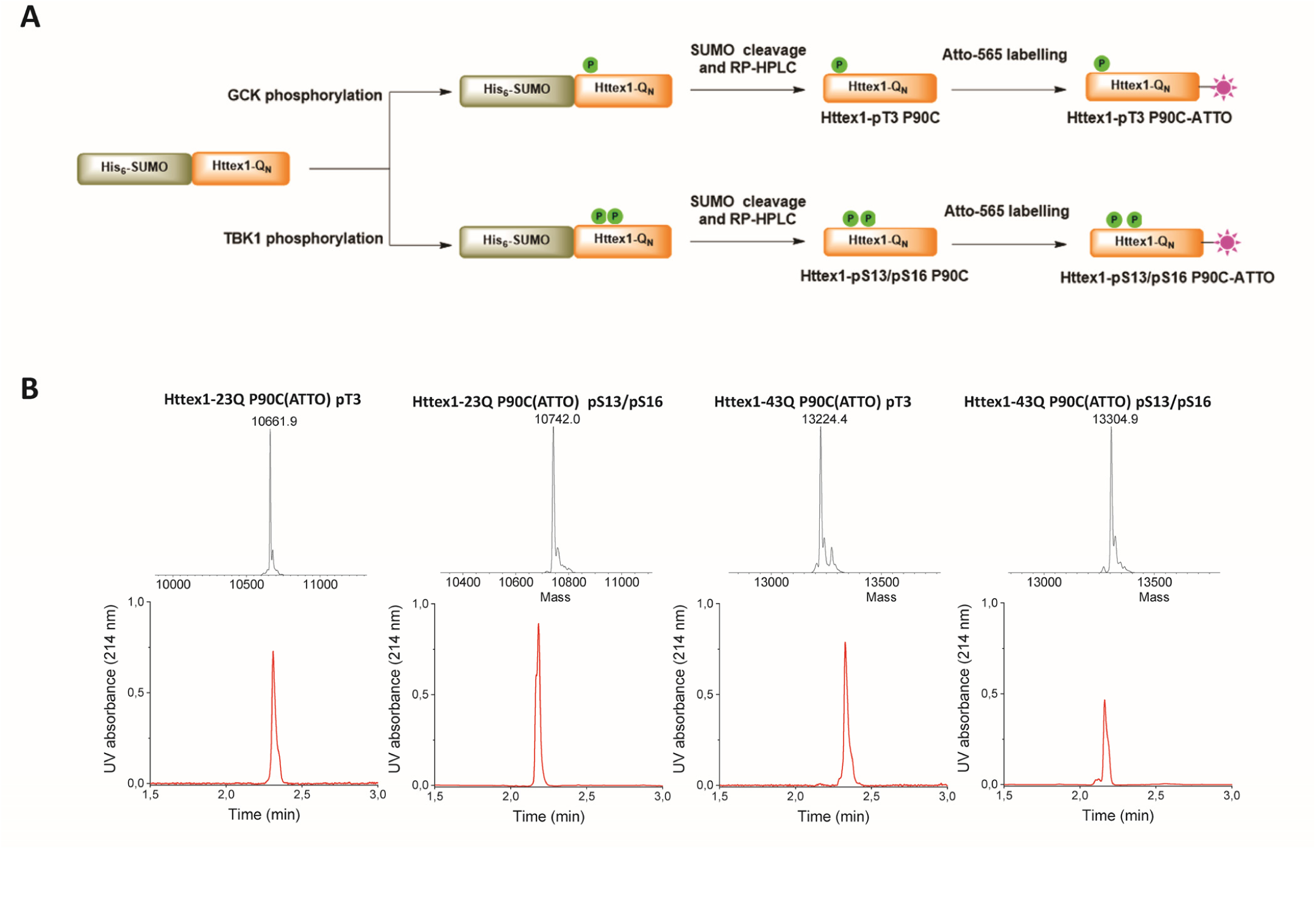
Atto-565-maleimide labeling of phosphorylated Httex1. **(A)** Schematic representation of the strategy to prepare and label Atto-565 phosphorylated Httex1. **(B)** Final characterization by ESI/MS and UPLC of the indicated labeled phosphorylated Httex1 proteins. (Expected molecular weights for the proteins are listed in **Table S1**).

**Figure 5.**
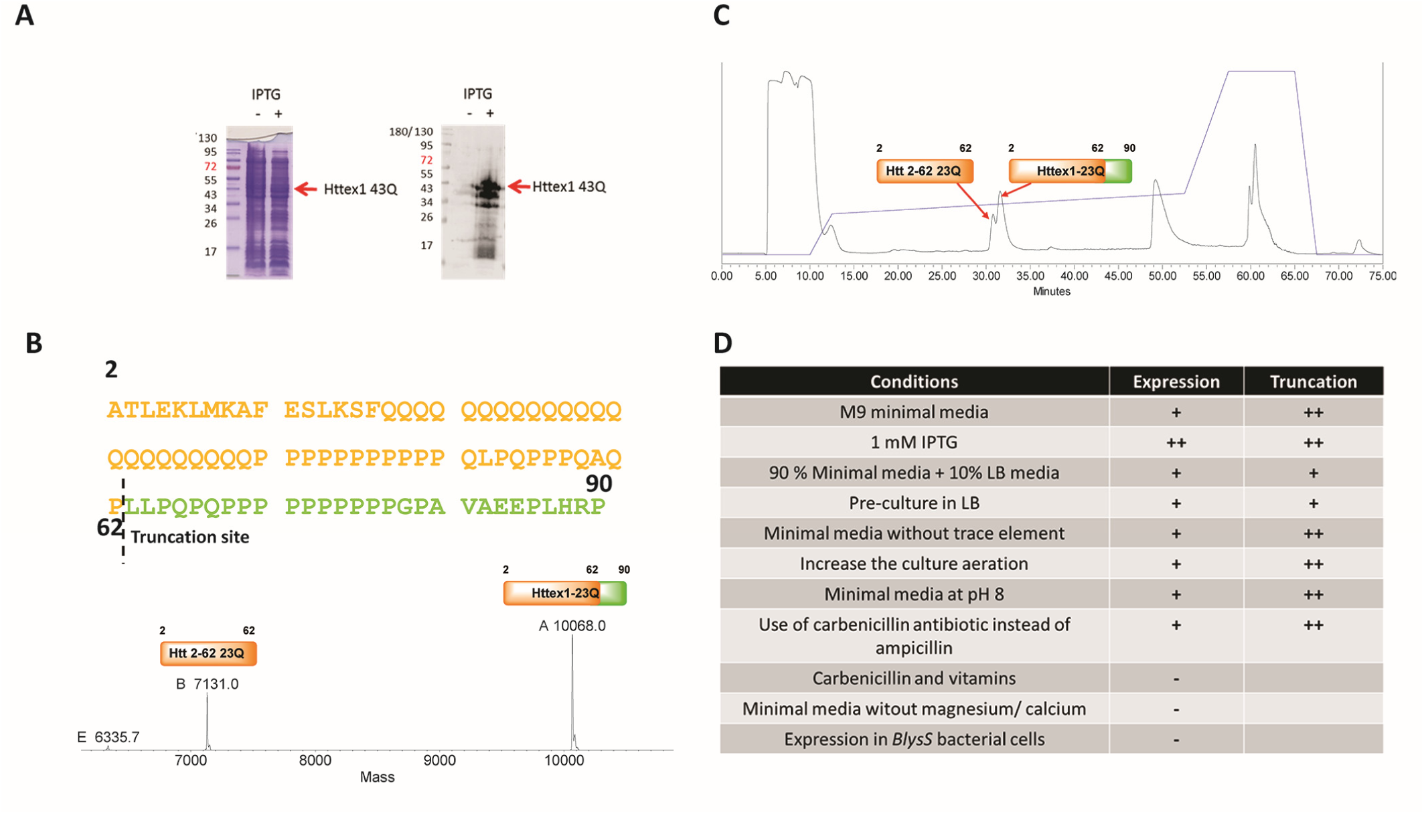
Optimizing the expression of SUMO-Httex1-Qn in minimal media. **(A)** SDS-PAGE and western blot analysis of the expression of SUMO-Httex1-43Q, highlighting the increased levels of truncation. **(B)** Analysis by ESI/MS of the elution fractions after HPLC of Httex1-23Q ^15^N. The truncation position is highlighted within the sequence of Httex1-23Q (upper panel). **(C)** HPLC chromatogram showing the purification of Httex1-23Q ^15^N after SUMO cleavage highlighting the HTT 2-62 truncation. **(D)** List of all the conditions that were tested to avoid truncation or lack of expression and obtained results (**+** for high, **++** for very high, and **–** for absence).

Using the protocol described above, we produced phosphorylated Httex1-Qn P90C (Httex1-23Q pT3 P90C, Httex1-43Q pT3 P90C, Httex1-23Q pS13/pS16 P90C and Httex1-43Q pS13/pS16 P90C) (Figure S5 and Figure S6- A). Then, to perform the labeling, these proteins were disaggregated using neat TFA, as described by Reif et al. [41]. Next, the thin protein film was resuspended in 100 mM Tris pH 7.4, 6 M GdHCl, 50 mM trehalose, 0.5 M proline, and 1.1 equivalents TCEP [56], and the pH was adjusted to 7.4. The fluorescent dye Atto-565-maleimide (1.5 equivalents) was added to the protein, and the reaction was kept on ice for 30 min. The labeling reaction was monitored by ESI/MS, with the knowledge that the addition of ATTO-565 to a protein corresponds to the addition of 633 Da to the molecular mass. All 4 phosphorylated proteins were successfully fluorescently labeled as evidenced by the appearance of an additional 633 Da to their molecular weight, as demonstrated by ESI/MS (Figure S6-B). After removal of the excess dye using a PD-10 column, the labeled phosphorylated proteins were purified by RP-HPLC, and the fractions containing the proteins were pooled and lyophilized. Figure 4-B shows the final characterization of Httex1-23Q pT3 P90C, Httex1-43Q pT3 P90C, Httex1-23Q pS13/pS16 P90C and Httex1- 43Q pS13/pS16 P90C by ESI/MS and UPLC.

### Generation of phosphorylated ^13^C- and ^15^N-labeled Httex1 suitable for NMR applications

Solution NMR spectroscopy has provided valuable insights into the structure and dynamics of soluble N-terminal peptides and Httex1 constructs in several works [57-60]. Previously, isotopic labeling of phosphorylated Httex1 was unaffordable by chemical synthesis, given the high costs of isotope-enriched amino acids. Having identified several kinases that phosphorylate Nt17 and developed an efficient strategy for the generation of recombinant Httex1 [41], we next sought to combine these advance to produce phosphorylated and isotopically labeled Httex1. When SUMO-Httex1-Qn was expressed in minimal media containing isotopically labeled ammonium chloride (^15^N) and glucose (^13^C), as previously reported [61], we observed an increase in to cleavages in the sequence of the fusion protein (Figure 5-A) leading to the generation of a fragment with a molecular weight of 7131 Da that corresponds to HTT 2-62, as indicated by ESI/MS in Figure 5-B. This fragment was challenging to separate from Httex1 by HPLC (Figure 5-C); the peaks of the 2 fragments did not resolve despite different gradient optimizations. Different conditions and media were tested (Figure 5-D), but we observed significant truncations or a lack of protein expression under all conditions. The best alternative method we identified was to grow the bacterial cells in standard LB media, and when the optical density of the culture reached 0.3, the cells were centrifuged and transferred to the isotopically labeled minimal media followed by induction of the expression with IPTG overnight at 18°C was performed [62]. Using this protocol, we succeeded in expressing and purifying the doubly labeled WT or mutant SUMO-Httex1 through nickel affinity chromatography (Figure S7-A and S7-B), which was then subjected to *in vitro* phosphorylation by GCK or TBK1 (Figure S7-A and S7-B) followed by ESI/MS (Figure 6-A and 6-B). We showed, as expected, that GCK phosphorylated SUMO-Httex1-23Q and Httex1-43Q at a single site (pT3) (Figure 6-A) and TBK1 phosphorylated at 2 sites (pS13/pS16) (Figure 6-B). The SUMO cleavage of the phosphorylated SUMO-Httex1-Qn ^13^C/^15^N was monitored by UPLC (Figure S7-C), and complete cleavage was indicated by the disappearance of the fusion protein peak and the appearance of 2 peaks corresponding to phosphorylated Httex1 and the SUMO tag (Figure S7-C). Removal of the SUMO tag and purification of the labeled proteins by RP-HPLC was performed (Figure S7-D). The purity of the proteins was verified by ESI/MS, UPLC, and SDS-PAGE (Figure 6- C).

**Figure 6.**
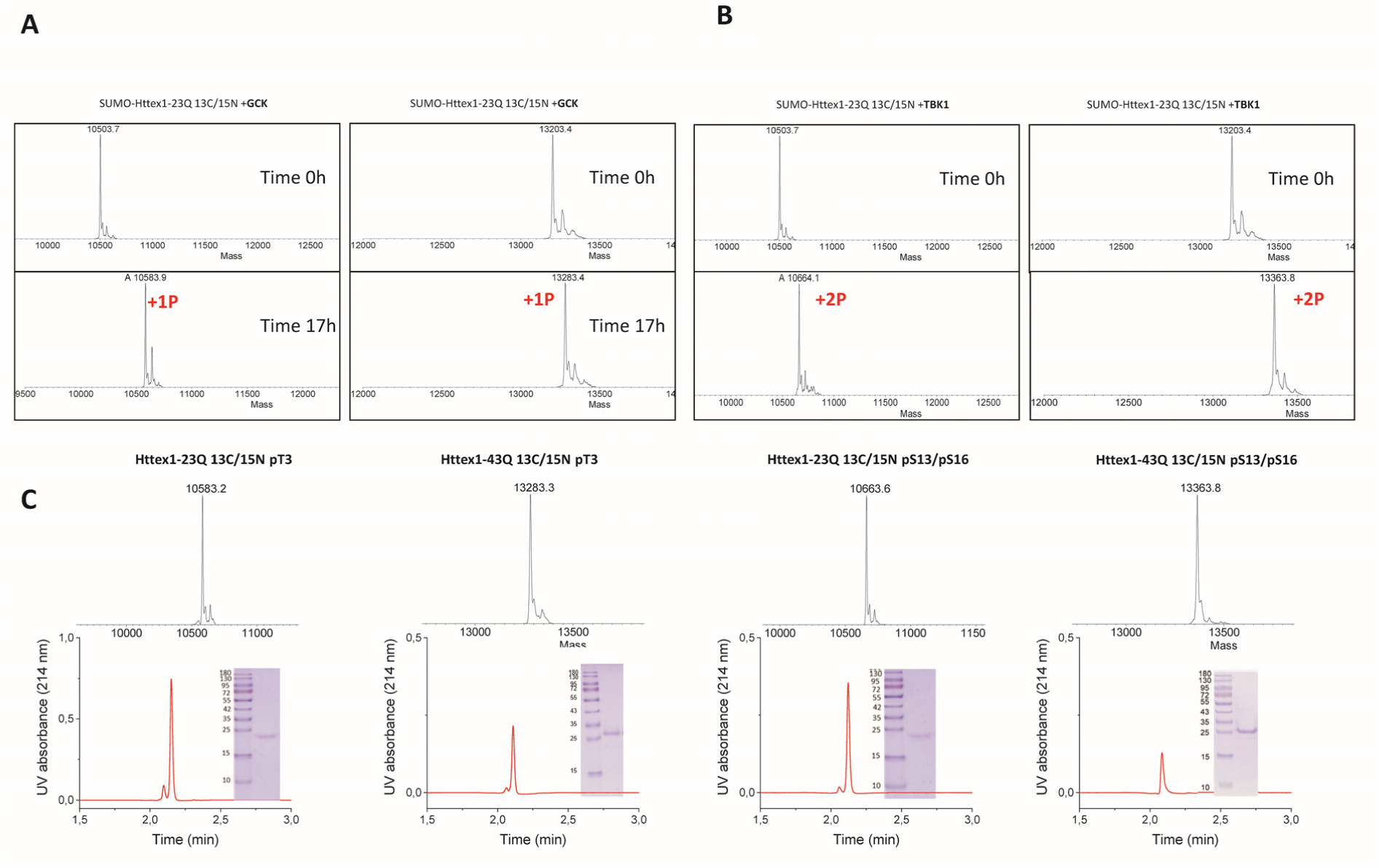
Phosphorylation and characterization of ^13^C/^15^N phosphorylated Httex1.**(A)** Monitoring of the phosphorylation reaction of SUMO-Httex1-23Q ^13^C/^15^N and SUMO-Httex1-43Q ^13^C/^15^N by GCK. **(B)** Monitoring of the phosphorylation reaction of SUMO-Httex1-23Q ^13^C/^15^N and SUMO-Httex1-43Q ^13^C/^15^N by TBK1 after an analytical SUMO cleavage. **(C)** Final characterization by ESI/MS, UPLC, and SDS-PAGE of Httex1-23Q ^13^C/^15^N pT3, Httex1-43Q ^13^C/^15^N pT3, Httex1-23Q ^13^C/^15^N pS13/pS16, and Httex1-43Q ^13^C/^15^N pS13/pS16. (Expected molecular weights for the proteins are listed in **Table S1**).

### Structural NMR studies of Httex1-23Q pT3 and Httex1-23Q pS13/pS16

With these proteins in hand, we then sought to gain insight into effect of Nt17 phosphorylation on the structural properties of Httex1. First, to test the quality and utility of our ^15^N/^13^C labeled samples, we verified that ^1^H spectra did not contain substantial amounts of unlabeled species, thus validating the success of the expression strategy presented above. ^1^H-^15^N HSQC NMR spectra were easily attainable for all the different proteins (unmodified, Httex1-23Q-pT3 and Httex1-23Q-pS13pS16) (Figure 7-A and 7-B). The signal spread, linewidths and stability over the course of one week for both phosphorylated forms were similar to those of the unmodified protein, indicating preserved disorder and equal or slower aggregation rates (Figure 7-A and 7-B).

**Figure 7.**
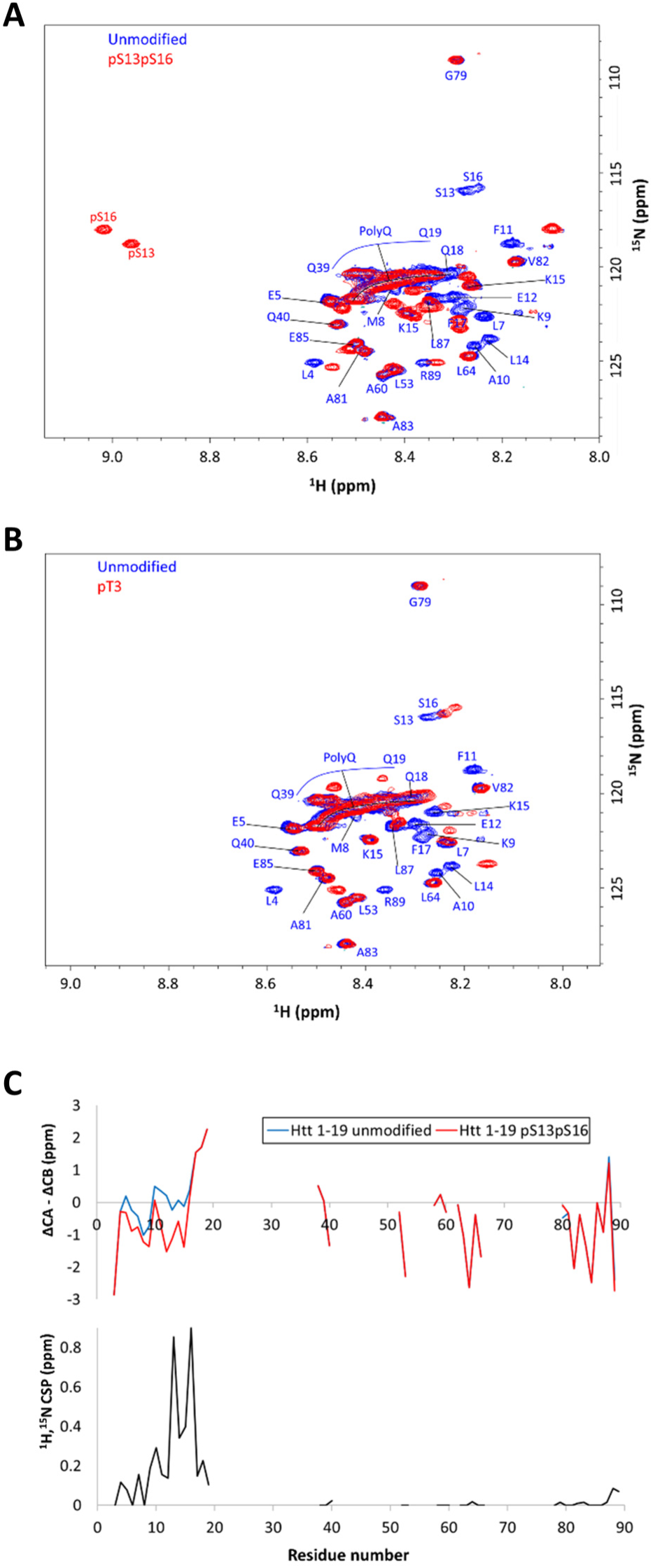
^1^H-^15^N HSQC NMR spectra of **(A)** Httex1-23Q pS13/pS16 and **(B)** Httex1-23Q pT3 in red, compared to the HSQC spectrum of unmodified Httex1-23Q in blue. All spectra were acquired at pH 7 and 25°C. The most important assignments of the unmodified and pS13pS16 forms are depicted. **(C)** The weighted chemical shift perturbations in ^1^H and ^15^N between the unmodified and pS13pS16 proteins (bottom) and a proxy for secondary structure propensities calculated as the difference of the residual CA and CB chemical shifts after subtraction of the random coil values (top). Notice that the slight beta propensity around residues 12 and 15 of Httex1-23Q pS13/pS16 could well arise as an artifact induced by the strong negative charges on Ser13 and Ser16 upon the subtraction of the random coil shift, without actually having any real structural implication (for pS13 and pS16 themselves we subtracted random coiled shifts of the phosphorylated amino acids, alleviating this problem).

After assigning the backbone resonances of our unmodified Httex1 construct in the conditions of interest (pH 7, 25°C, Table S2), we could use this information to interpret the differences between the HSQC spectra of unmodified and both phosphorylated forms. For Httex1-23Q ^13^C/^15^N pS13/pS16 (Figure 7-A), we observed strong downfield perturbations for S13 and S16 chemical shifts in the ^1^H dimension, as documented for phosphorylated serine [63, 64], along with smaller but sizeable perturbations of other residues within the Nt17 region including the first two glutamines of the polyQ tract (Q18 and Q19). However, we did not observe significant perturbations for the terminal glutamines (Q39 and Q40), nor for residues in the proline-rich or C-terminal regions. Similarly, for Httex1 ^13^C/^15^N pT3 (Figure 7-B), we observed small but sizeable effects on the Nt17 residues, but not for the residues of the proline-rich and C-terminal regions. It is to be noted that no crosspeak for phosphorylated T3 was visible at pH 7 and 25°C. Moreover, T3 is also undetected in the unmodified protein under these conditions of neutral pH; however, we observed at pH 4.2 or lower a crosspeak at chemical shifts compatible with phosphorylated threonine while the crosspeaks for S13 and S16 remained close to their positions in the HSQC spectrum of the unmodified protein (Figure S8-B).

Next, we obtained the backbone resonance assignments of the Httex1 construct doubly phosphorylated at S13 and S16, enabling a detailed residue-wise comparison of chemical shifts in contrast to the unmodified protein (Figure 7-C and Table S2). The phosphorylated serines exhibited large ^1^H,^15^N perturbations, as expected, and were the largest. Away from the serine residues, the ^1^H,^15^N chemical shift perturbations slowly decay over the sequence but peaking at residues 4, 7, 10, and 18. The location of these residues relative to S13 and S16 follows closely the spacing expected in a helical structure. Notably, in the unmodified protein, the ^13^C chemical shifts of CA and CB atoms indicated that this protein exhibited a large disorder but with a substantial helical propensity, especially at residues 17-19, consistent with previous reports [58, 60]. This helical propensity is retained in Httex1-23Q ^13^C/^15^N pS13/pS16 according to the CA and CB chemical shifts (Figure 7-C). To further explore the weak alpha-helical propensities, we titrated solutions of ^15^N labeled proteins with trifluoroethanol (TFE), an agent well-known to expose helical structures. This experiment revealed large chemical shift dispersions on the unmodified and both phosphorylated forms, suggesting a similar latent helical propensity in all forms (Figure S8-A).

The pH of the buffer could also potentially affect the structure; therefore, we assessed the phosphorylated and unmodified proteins at different pHs. Increasing the pH above neutrality caused most of the signals to rapidly broaden beyond detection, starting at pH 7.5, in all the proteins (Figure S8-C). On the other hand, when decreasing towards acidic pH, we found only slight shifts in the unmodified protein, suggesting virtually no substantial effects on the structure and dynamics pf the proteins (Figure S8-C). In the phosphorylated forms, the crosspeaks for pS13 and pS16 shifted considerably upfield as the pH decreased, most likely reflecting just protonation of the phosphate groups (Figure S8-C). At the same time, crosspeaks of all the other residues were only slightly affected, indicating no substantial effects of the pH on the structure and dynamics of proteins. In all forms, a crosspeak likely corresponding to T3 appeared at pH 4.2 or lower, but this only reflects the slower H-N exchange at low pH without any significant effects on the other residues, nor does have any structural implications. In all cases, individual residues such as His88, Glu12, and sequence neighbors experienced shifts directly related to protonation as the pH decreased, but, again, these were likely just their protonation and not structural effects (Figure S8-C).

### Real-time monitoring of TBK1 phosphorylation of Httex1 suggests that S13 phosphorylation occurs first and primes phosphorylation at S16

The high sensitivity of NMR spectroscopy to phosphorylation and its simplicity, pose it as a valuable tool for real-time monitoring of Httex1 phosphorylation by kinases. Therefore, we sought to use to assess potential cross-talk between phosphorylation at S13 and S16, as suggested by previous studies [29]. This experimental setup is much more straightforward than the use of antibody-based approaches such, has exquisite time resolution, and provides residue-level information on the structural consequences of phosphorylation, if any. We subjected the doubly labeled Httex1-23Q at a concentration of 200 µM to phosphorylation by TBK1 in a compatible buffer with excess Mg-ATP (Figure 8). The reaction was performed at 20°C, in order to slow the enzymatic reaction adequately and monitor it in real-time by HSQC spectra collected in under 3 minutes each, at high resolution without distortions. We observed the rapid decay of the crosspeaks corresponding to both serine residues (Figure 8). In parallel, a new crosspeak appeared close to them, and another appeared in a downfield region consistent with a phosphorylated serine. Over-time, these two crosspeaks, tentatively assigned to an intermediate, faded out as those assigned to pS13 and pS16 in the pure pS13pS16 protein appeared and gained intensity (Figure 8). A ^1^H,^13^C plane of a CBCA(CO)NH experiment collected 21 hours after initiation of the reaction revealed that the crosspeak for a phosphorylated serine in the intermediate shares ^13^C chemical shifts with Ser13 in the pS13pS16 protein (Figure S9). We concluded that the intermediate corresponds to a species where Ser13 is phosphorylated but Ser16 is not (S16 gives its crosspeak in the spectral region of unphosphorylated serines). Interestingly, this was confirmed by the absence of a fourth cross-peak in the region of phosphorylated serines that would have been attributed to an intermediate phosphorylated at S16 with S13 unmodified. These observations suggest that S13 phosphorylation primes phosphorylation at S16, similar to what has been suggested previously [29, 65]. Additionally, these findings were in accordance with our previous cellular and *in vivo* observation where we showed that TBK1 phosphorylated mainly S13 [43].

**Figure 8.**
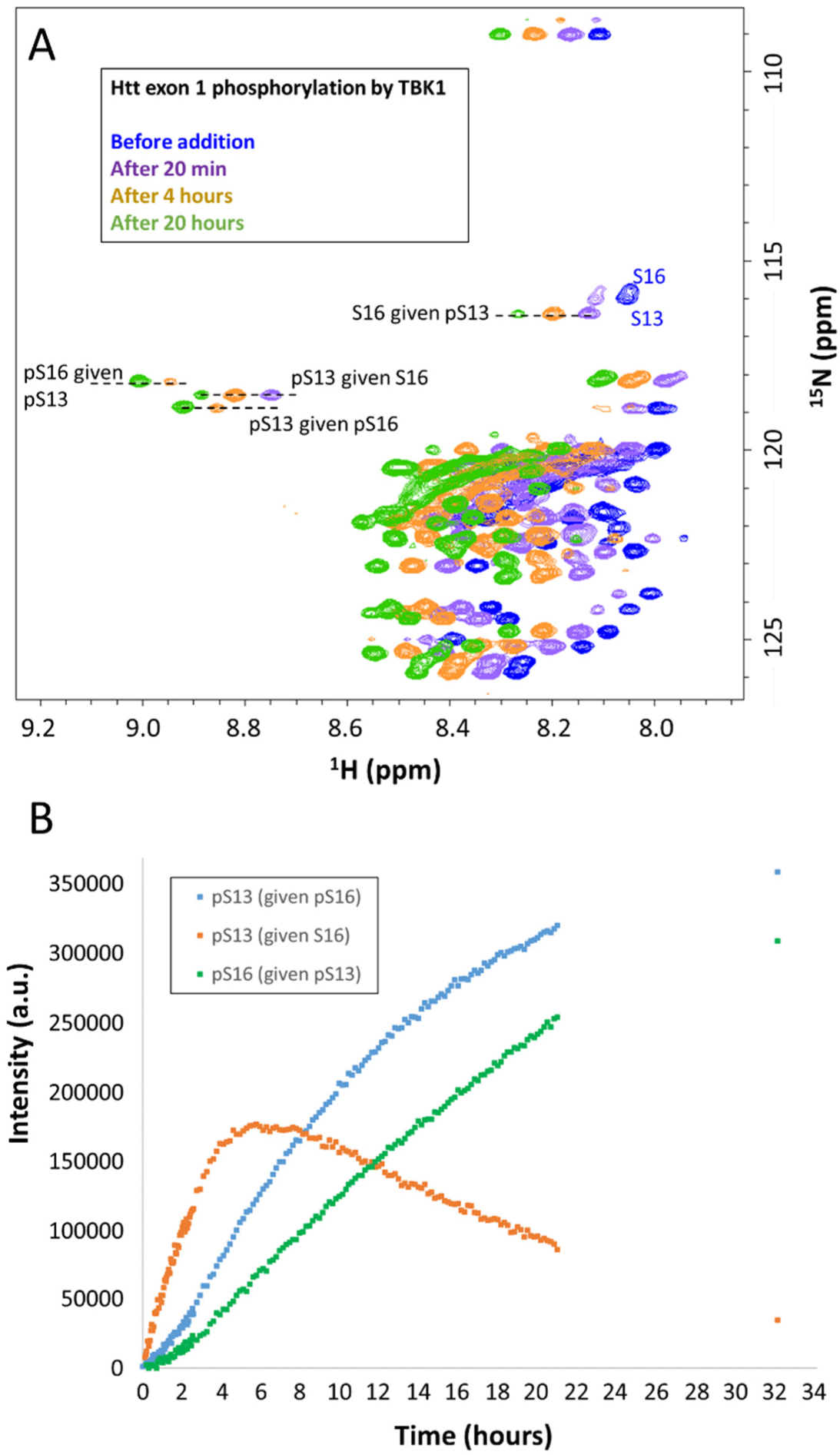
Phosphorylation of Httex1-23Q 13C/15N followed by NMR. Monitoring of 500 µl of a 200 µM sample of doubly labeled Httex1-23Q as it is phosphorylated by TBK1 in buffer containing 15 μg of TBK1 plus 5 mM Mg-ATP and a suitable buffer (30 mM HEPES, 5 mM MgCl_2_, 5 mM EGTA, 1 mM DTT, pH 7.2) at 20°C. Each HSQC spectrum was acquired in 2 min and 50 seconds with a 0.5 recycle delay and 128 ^15^N increments. Spectra were acquired serially over 21 hours with one final acquisition at 32 hours. **(A)** An example spectrum taken before TBK1 addition and after 20 min, 4 hours and 21 hours, offset to the left at increasing times for clarity; assignments of all observed S13 and S16 species are indicated. **(B)** The intensities of these cross-peaks over time, stressing the initial phosphorylation of S13 with subsequent phosphorylation at S16 leading to the final pS13/pS16 product.

## Conclusions

We have described here the first method for enzymatic generation of site-specifically phosphorylated WT and mutant Httex1 proteins. This was enabled by 1) the discovery and validation of novel kinases that efficiently phosphorylate Httex1 at S13 and S16 (TBK1[43]), T3 (GCK) or T3 and S13 (TNIK and HGK); 2) developing conditions that allow performing *in vitro* phosphorylation reactions on mutant HTT under conditions where the protein is stable and remains soluble (SUMO-fusion proteins); and 3) developing efficient methods for the cleavage and purification of the final phosphorylated proteins. The use of the newly discovered kinases combined with the use of the SUMO fusion protein expression and purification method for producing recombinant Httex1 allowed the production of milligram quantities of site-specifically phosphorylated Httex1 proteins.

As a proof of concept, we used GCK- and TBK1-mediated phosphorylation protocols to produce unmodified or site-specific fluorescently labeled WT and mutant Httex1 phosphorylated at T3 or S13/S16. This approach could replace existing strategies based on the use of large fusion fluorescent proteins or the nonspecific introduction of fluorescent dyes onto lysine residues. In addition, we showed that our new method also addresses previous challenges associated with the generation of site-specifically phosphorylated and isotopically labeled mutant Httex1 proteins for structural studies by solution and solid-state NMR [46, 47]. Previous NMR studies have focused primarily only on investigating the structure of unmodified monomeric [57] and fibrillar [47] Httex1 proteins. Herein, we developed an optimized expression protocol for Httex1 in minimal media, which significantly reduces non-desired cleavages, thus enhancing the yield of recombinant isotopically labeled untagged, native or phosphorylated Httex1 suitable for NMR studies (Figure 6). We showed that phosphorylation of the unmodified protein could be easily monitored by NMR spectroscopy with high structural and time resolution (Figure 8). We also demonstrated that pT3 pS13/pS16 Httex1 remain largely disordered, with a gradient of helical propensity that starts around the first glutamine residues of the polyglutamine tract and gradually fades downstream, as in the unmodified protein. Our preliminary studies show that phosphorylation affects the chemical shifts of the target residues and their neighboring residues but mostly through electrostatic effects on the chemical shifts themselves and without any substantial effect on either the structure or dynamics. In particular, the last glutamine residues of the polyQ tract and all downstream residues through to the end of Httex1 appear to be insensitive to phosphorylation. Furthermore, we establish for the first time that phosphorylation at S13 could prime phosphorylation at neighboring S16, thus underscoring the importance of further studies to investigate cross-talk between different Nt17 PTMs.

This new method described here eliminates the need to use phosphomimetic mutations at these residues and addresses many of the limitations of existing protein semisynthetic strategies for producing these proteins, including the requirement to introduce a nonnative residues at residue 18 (Q18A) [66]. Furthermore, unlike protein semisynthesis, this method is affordable, accessible, and enables the generation of milligram quantities of highly pure phosphorylated proteins. Finally, the high solubility of the SUMO-Httex1 fusion protein enables greater flexibility to handle and manipulate the protein, thus facilitating the introduction of PTMs and chemical functional groups or probes (e.g., a fluorescent dye, biotin) into mutant Httex1, which was not feasible in the absence or after removal of the SUMO tag.

We believe that these advances will open new opportunities and expand the experimental approaches aimed at elucidating the role of PTMs in regulating HTT structure, aggregation, pathology formation, and cell-to-cell propagation. Moreover, the method described here can be extended to other PTMs, such as ubiquitination, SUMOylation, and acetylation, once the enzymes involved in regulating these modifications are identified. Finally, the ability to generate homogeneously modified proteins should facilitate the development of novel assays to quantify the levels of Nt17 modified forms of HTT in biological samples and assess their potential as biomarkers for early diagnosis or to monitor HD progression.

## Experimental Section

### Materials

The pTWIN1 vector, containing human Httex1 fused to His6-SUMO, was ordered from GeneArt Gene Synthesis (Regensburg-DE). *E. coli* B ER2566 was from New England Biolabs (Ipswich-USA). Ampicillin, DTT, isopropyl β-D-1- thiogalactopyranoside (IPTG), and PMSF were obtained from AppliChem. Imidazole, cOmplete Protease Inhibitor Cocktail, magnesium chloride (MgCl_2_), magnesium sulfate (MgSO_4_), and trifluoroacetic acid were purchased from Sigma-Aldrich Chemie GmbH, (Buchs-CH). EGTA solution was from Boston Bioproducts (Chestnut-USA). Mg-ATP was from Cayman chemical (Ann Arbor-USA). EDTA was from Fisher Scientific (Ecublens-CH), and Luria Broth (Miller’s LB Broth) was from Chemie Brunschwig (Basel-CH). HPLC-grade acetonitrile was from Macherey-Nagel (Oensingen-CH). The spectrophotometer semimicro cuvette was from Reactolab (Servion-CH). The C4 HPLC column was from Phenomenex (Basel-CH). The HisPrep 16/10 column was from GE healthcare (Dietikon-CH). Purified recombinant TBK1 (0.62 mg/µl, cat. # NM_013254) and GCK kinases (0.5 mg/µl, cat. # BC047865), from MRC PPU Reagents and service, Dundee-SCO, were stored at -80°C. Rabbit antibodies against pT3, pS13, and pS16 were homemade.

### Instruments

A Vibra-cell VCX130 ultrasonic liquid processor (Sonics), an Äkta 900 equipped with a fraction collector (GE Healthcare), and Waters UPLC and HPLC systems were used. For ESI-MS, a Finnigan LTQ (Thermo Fisher Scientific) was employed. A lyophilizer instrument (FreeZone 2.5 Plus) and a shaking incubator with a temperature regulator (Infors HT multitron Standard) were also used in the experiments.

### Kinase screening (IKPT service)

Kinase screening (IKPT service) was performed by Kinexus, Canada. Httex1-23Q at 1.4 ⍰M and WT Nt17 peptide (ATLEKLMKAFESLKSF) at 150 ⍰M was screened against a panel of 298 selected Ser/Thr kinases using a radiometric assay method with [γ-^33^P]ATP. The assay was initiated by the addition of [γ33P] ATP and the reaction mixture incubated at ambient temperature for 30 minutes. After the incubation period, the assay was terminated by spotting 10 µl of the reaction mixture onto a multiscreen phosphocellulose P81 plate. The multiscreen phosphocellulose P81 plate was washed 3 times for approximately 15 minutes each in a 1% phosphoric acid solution. The radioactivity on the P81 plate was counted in the presence of scintillation fluid in a Trilux scintillation counter.

### Expression of His6-SUMO-Httex1-Qn (n= 23 or 43)

The expression of His-SUMO-Httex1-Qn was performed as previously reported [41]. A pTWIN1 plasmid containing SUMO-Httex1-Qn with ampicillin (Amp) resistance was transformed into *E. coli* ER2566 using the heat shock method [67], and then the transformed bacteria were plated in an agar plate with ampicillin resistance. Next, a single colony was inoculated in a culture flask with 400 ml of LB+Amp (100 µg/ml) medium and incubated with shaking (190 rpm) at 37°C overnight (preculture). Large expression of the protein (12 L) started the day after, at an optical density (OD_600_) of 0.15 was observed in multiple 5 L flasks (3 L of culture maximum per flask). When the OD_600_ reached 0.5 to 0.6, the expression of HIS-SUMO-Httex1-Qn was induced with IPTG at a final concentration of 0.4 mM, and the culture was incubated at 18°C for 18 hours (overnight). The cells were harvested by centrifugation (3993 × g, 4°C, 10 min), and the cell pellet was kept on ice for further purification.

### Expression of His6-SUMO-Httex1-Qn (n= 23 or 43) in minimal isotopic media

The transformation of the plasmid and the preculture were performed, as mentioned above in LB+Amp media. Then, a large culture was grown first in LB+Amp media as similarly described to an OD_600_ of 0.15. When the OD_600_ reached 0.3, the cells were centrifuged (3993 × g, 4°C, 10 min) and resuspended in minimal media containing ^15^N-labeled ammonium chloride and ^13^C-labeled glucose for the production of ^13^C/^15^N-labeled His-SUMO-Httex1-Qn. Then, the culture was induced at an OD less than 0.6 and incubated overnight at 18°C. The cells were harvested by centrifugation (3993 × g, 4°C, 10 min), and the cell pellet was kept on ice for immediate purification.

### Immobilized metal affinity chromatography (IMAC) purification

The bacterial pellet was resuspended in IMAC buffer A (50 mM Tris, 500 mM NaCl, 30 mM imidazole, pH 7.5, filtered through a 0.65 µm filter) supplemented with PMSF and complete protease inhibitors and then sonicated on ice for cell lysis (70% amplitude, total sonication time of 5 min, intervals of 30 seconds of sonication followed by a 30 seconds pause). The cell lysate was centrifuged (39191 × g, 4°C, 60 min), and the supernatant was filtered (0.45 µm, syringe filters) and loaded onto the Ni-NTA column on fast-performance liquid chromatography (FPLC) system at 4°C. The protein was then eluted with 100% IMAC buffer B (50 mM Tris, 500 mM NaCl, 500 mM imidazole, pH 7,5, filtered through a 0.65 µm filter). Coomassie SDS-PAGE was used to analyze the eluted fractions, and the fractions containing SUMO-Httex1-Qn were pooled together and kept on ice for subsequent phosphorylation.

### Quantitative in vitro phosphorylation of SUMO-Httex1-Qn by GCK and TBK1

SUMO-Httex1-Qn after IMAC purification was dialyzed against 4 L of TBS buffer overnight at 4°C, and its concentration was measured using a nanodrop UV spectrophotometer. The concentration was calculated by measuring the absorbance of the dialyzed or desalted protein solution at 280 nm, where the extinction coefficient was 1490 M-1 cm-1. Ten milligrams of the fusion protein was removed, and the volume was adjusted to 18 mL. Then, 2 mL of 10X phosphorylation buffer (50 mM MgCl_2_, 80 mM EGTA, 40 mM EDTA, 10 mM DTT) was added. The pH of the mixture was adjusted to 7.4, and Mg-ATP was added at a final concentration of 5 mM. Finally, GCK or TBK1 was added at a ratio of 1:30 w/w to Httex1-Qn (666 µL in the case of GCK and 537 µL in the case of TBK1), and the enzymatic reaction was incubated at 30°C overnight (17 hours). To determine the completion of phosphorylation, 30 µL was removed and supplemented with 1 µL of ULP1 (1 mg/mL) to cleave the SUMO tag for analytical analysis. Ten microliters were injected for analysis by LC-ESI-MS (positive ionization mode). When the ESI/MS confirmed complete phosphorylation, the reaction solution was kept on ice for further processing.

### SUMO cleavage and HPLC purification

To cleave the SUMO tag from Httex1-Qn, 0.6 mL of ULP1 enzyme was added to the phosphorylation reaction mixture and incubated on ice for 15 min. Once UPLC confirmed SUMO cleavage, phosphorylated Httex1-Qn was purified by HPLC on a C4 column using a gradient of 25%-35% solvent B (HPLC-grade acetonitrile containing 0.1% v/v TFA) in solvent A (ultrapure water containing 0.1% v/v TFA). Collected fractions were analyzed for the presence of the desired protein by ESI/MS and pooled accordingly for lyophilization. The purity of the lyophilized protein was assessed by ESI/MS (ESI/MS spectra were deconvoluted with MagTran software), UPLC, and SDS-PAGE.

### Fluorescent labeling with Atto-565-maleimide

One milligram of phosphorylated Httex1-Qn P90C was disaggregated using neat TFA, as described by Reif et al. [41]. Then, the thin protein film was resuspended in 100 mM Tris, pH 7.4, 6 M GdHCl, 50 mM trehalose, 0.5 M proline, and 1.1 equivalents of TCEP [56], and the pH was adjusted to 7.4. Then, 1.5 equivalents of the fluorescent dye Atto-565-maleimide was added to the protein, and the reaction was kept on ice for 30 min. The reaction was monitored by ESI/MS, and upon completion, excess Atto-565-maleimide was removed using a PD10 column equilibrated with 20% acetonitrile in water. The protein was immediately injected onto a C4 300 Å 250 × 4.6 mm column. The protein was eluted using a gradient of 25 to 55% solvent B over 50 min. The collected fractions were analyzed by ESI/MS for the presence of the proteins of interest, and the corresponding fractions were pooled according to the different qualities. The final purity of the protein was determined by UPLC and ESI/MS.

### Nuclear magnetic resonance (NMR) spectroscopy

All NMR experiments were carried out in a Bruker Avance III 800 MHz spectrometer equipped with a CPTC cryoprobe. Spectra for protein analysis and resonance assignments were collected at 25°C on protein samples with approximately 100 µM concentrations, while spectra for monitoring phosphorylation kinetics were collected on 200 µM protein samples to achieve strong sensitivity in short acquisition times and at 20°C to slow the kinetics. Samples for phosphorylation kinetics were prepared in 30 mM HEPES, 5 mM MgCl_2_, 5 mM EGTA, 1 mM DTT, and 5 mM Mg-ATP with 10% D_2_O at pH 7.2 adjusted after the addition of ATP and prior to the addition of 15 µg of TBK1. All other samples were prepared in 10 mM Na_2_HPO_4_ and 10% D_2_O buffer at pH 7. All spectra were acquired and processed using Bruker TopSpin 4.0 and analyzed with CARA and Sparky-NMRFAM.

HSQC spectra for monitoring the phosphorylation kinetics were acquired using a sensitivity-enhanced sequence with 2 scans, a short recycle delay of 0.5 seconds and 128 ^15^N increments, (processed with 256 indirect points), a combination that in our hands resulted in high resolution, excellent intensity and no distortions in 2 min and 50 seconds of total acquisition time, which is relatively short compared to the phosphorylation rate in the set conditions. Resonance assignments for the unmodified and pS13pS16 proteins were obtained by analyzing a high-resolution HSQC experiment (sensitivity enhanced, 256 indirect points, 1 second recycle delay) and standard triple-resonance experiments: HNCO, HN(CA)CO, HNCA, HN(CO)CA, CBCA(CO)NH and HNCACB. All 3D experiments were acquired using standard pulse sequences, with 40 increments in ^15^N, 128 increments in the ^13^C dimensions, 1 second of recycle delay, and nonuniform sampling (NUS) at 50% for HNCACB or 25% for the other five spectra. Resonance assignments were aided by previously published assignments of Httex1[58-60].

The chemical shift perturbations in ^1^H and ^15^N between the unmodified and pS13pS16 proteins were computed as:

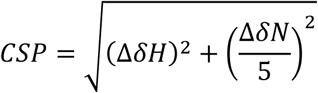

where ΔδH is the difference in ^1^H chemical shifts and ΔδN is the difference in ^15^N chemical shifts, and the factor 5 corresponds to the value widely used to weigh down the ^15^N shifts.

Secondary structure propensities from the ^13^C chemical shifts (ΔCA - ΔCB) were obtained by subtracting the random coiled CA and CB chemical shifts of the amino acids from the experimentally observed CA and CB shifts and then subtracting the two differences. This indicator is insensitive to offsets or calibration problems in the ^13^C dimension; a positive value above +1 indicates alpha helical propensity, and a negative value under -1 indicates beta sheet propensity. For phosphorylated serine, the CA and CB chemical shifts reported by [63] were used as random coil references; for the other amino acids, we used the random coil shifts from [68].

### Coexpression of Httex1-16Q-eGFP and kinases in HEK 293 cells

HEK 293 cells were cultured in 95% air and 5% CO2 in Dulbecco’s modified Eagle’s medium (Gibco) supplemented with 10% fetal bovine serum (Gibco) and penicillin-streptomycin (Thermo Fisher). Plasmids GCK (MSP4K2) (cat. no. RC200472, OriGene), HGK (MAP4K4) (cat. no. RC215163, OriGene), and TNIK (Plasmid #45276 Addgene) were acquired. Transfections were carried out by Lipofectamine 2000 according to the manufacturer’s protocol. Lysis was performed in RIPA lysis buffer (150 mM sodium chloride, Triton X-100, 0.5% sodium deoxycholate, 0.1% SDS (sodium dodecyl sulfate), 50 mM Tris, pH 8.0) supplemented with 1× protease and phosphatase inhibitor 2, 3 mixture (Sigma). Cell lysates were then centrifuged at 15000×g for 20 min, and the supernatant was collected as the soluble fraction. The protein concentration was measured using the BCA system, and approximately 20-60 µg of protein was processed for the WB assay. Antibodies used for western blotting were pT3-Huntingtin CHDI- 90001528-2 from CHDI/Thermo Scientific, pS13 Huntingtin CHDI-90001039-1 from CHDI/Thermo Scientific, pS16- Huntingtin in-house generated, and GAPDH/14C10) 2118S from Cell Signaling.

## Acknowledgements

This work was supported by funding from CHDI (R01NS086452). We are grateful to Rajasekhar Kolla for critical review of the manuscript and Nour Chiki-Benmrad for technical assistant during her semester project. We thank the MRC PPU Reagents and Services facility (MRC I PPU, College of Life Sciences, University of Dundee, Scotland, mrcppureagents.dundee.ac.uk) for providing TBK1 and GCK kinases on a cost-recovery basis.

## Graphic for Table of Contents

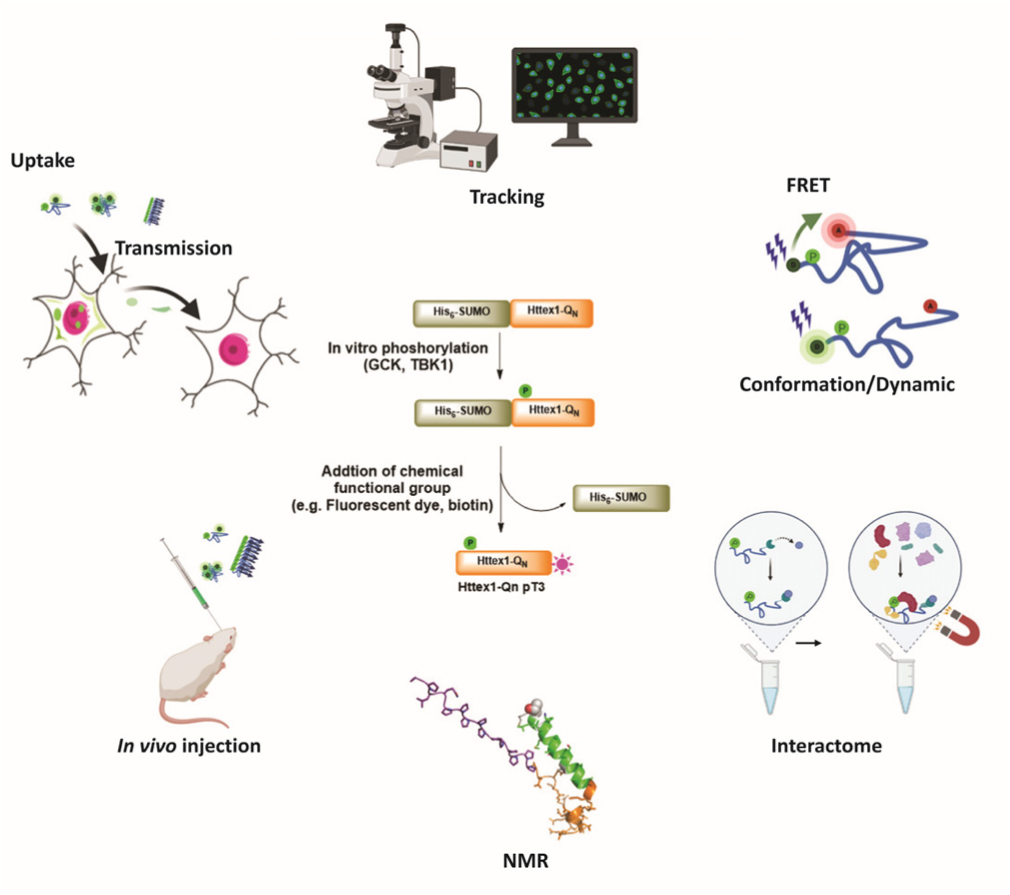

The tools presented in this paper offer new opportunities and enlarge the range of potential methods and experimental approaches that can be used to study the mechanisms by which the Nt17 phosphorylation influences the properties and functions of Httex1 in health and disease. (NMR depiction is adapted from [57]).

**Scheme 1.**
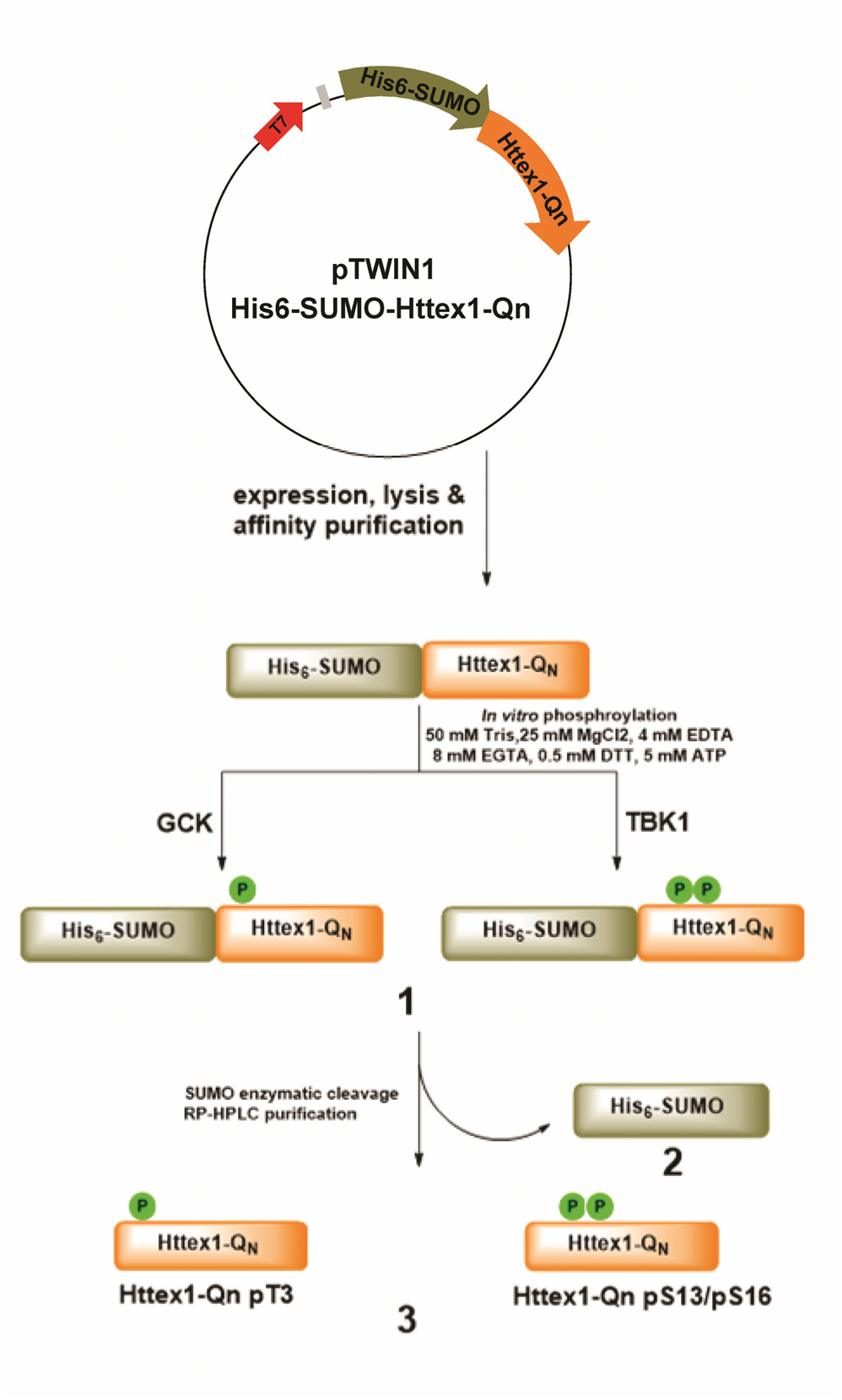
Schematic overview of the method used for the expression and purification of SUMO Httex1-Qn (n= 23 or 43), which undergoes *in vitro* phosphorylation by TBK1 or GCK to generate SUMO-Httex1-Qn pT3 or SUMOHttex1-Qn pS13/pS16 **(1**), then, the SUMO tag **(2)** was then cleaved by ULP1, and the desired phosphorylated Httex1-Qn **(3)** was purified by HPLC.

## References

1. Ross, C.A. and S.J. Tabrizi, Huntington’s disease: from molecular pathogenesis to clinical treatment. The Lancet Neurology, 2011. 10(1): p. 83–98.

2. Rosenblatt, A., et al., Predictors of neuropathological severity in 100 patients with Huntington’s disease. Annals of Neurology: Official Journal of the American Neurological Association and the Child Neurology Society, 2003. 54(4): p. 488–493.

3. Huntington, G., On chorea. 1872.

4. Kagel, M.C. and N.A. Leopold, Dysphagi in Huntington’s disease: A 16-year retrospective. Dysphagia, 1992. 7(2): p. 106–114.

5. Marder, K., et al., Rate of functional decline in Huntington’s disease. Neurology, 2000. 54(2): p. 452–452.

6. de Boo, G.M., et al., Early cognitive and motor symptoms in identified carriers of the gene for Huntington disease. Archives of Neurology, 1997. 54(11): p. 1353–1357.

7. Slaughter, J.R., M.P. Martens, and K.A. Slaughter, Depression and Huntington’s disease: prevalence, clinical manifestations, etiology, and treatment. CNS spectrums, 2001. 6(4): p. 306-308,325-326.

8. MacDonald, M.E., et al., A novel gene containing a trinucleotide repeat that is expanded and unstable on Huntington’s disease chromosomes. Cell, 1993. 72(6): p. 971–983.

9. Gusella, J.F., et al., A polymorphic DNA marker genetically linked to Huntington’s disease. Nature, 1983. 306(5940): p. 234–238.

10. Kremer, B., et al., A worldwide study of the Huntington’s disease mutation. The sensitivity and specificity of measuring CAG repeats. N Engl J Med, 1994. 330(20): p. 1401–6.

11. Snell, R.G., et al., Relationship between Trinucleotide Repeat Expansion and Phenotypic Variation in Huntingtons-Disease. Nature Genetics, 1993. 4(4): p. 393–397.

12. Morley, J.F., et al., The threshold for polyglutamine-expansion protein aggregation and cellular toxicity is dynamic and influenced by aging in Caenorhabditis elegans. Proceedings of the National Academy of Sciences, 2002. 99(16): p. 10417–10422.

13. Chen, S., F.A. Ferrone, and R. Wetzel, Huntington’s disease age-of-onset linked to polyglutamine aggregation nucleation. Proceedings of the National Academy of sciences, 2002. 99(18): p. 11884–11889.

14. Steffan, J.S., et al., The Huntington’s disease protein interacts with p53 and CREB-binding protein and represses transcription. Proceedings of the National Academy of Sciences, 2000. 97(12): p. 6763–6768.

15. Huang, C.C., et al., Amyloid formation by mutant huntingtin: threshold, progressivity and recruitment of normal polyglutamine proteins. Somatic cell and molecular genetics, 1998. 24(4): p. 217–233.

16. Waelter, S., et al., Accumulation of mutant huntingtin fragments in aggresome-like inclusion bodies as a result of insufficient protein degradation. Molecular biology of the cell, 2001. 12(5): p. 1393–1407.

17. Martindale, D., et al., Length of huntingtin and its polyglutamine tract influences localization and frequency of intracellular aggregates. Nature Genetics, 1998. 18(2): p. 150–154.

18. DiFiglia, M., Aggregation of Huntingtin in Neuronal Intranuclear Inclusions and Dystrophic Neurites in Brain. Science, 1997. 277(5334): p. 1990–1993.

19. Ratovitski, T., et al., Mutant Huntingtin N-terminal Fragments of Specific Size Mediate Aggregation and Toxicity in Neuronal Cells. Journal of Biological Chemistry, 2009. 284(16): p. 10855–10867.

20. Sathasivam, K., et al., Aberrant splicing of HTT generates the pathogenic exon 1 protein in Huntington disease. Proc Natl Acad Sci U S A, 2013. 110(6): p. 2366–70.

21. El-Daher, M.T., et al., Huntingtin proteolysis releases non-polyQ fragments that cause toxicity through dynamin 1 dysregulation. Embo Journal, 2015. 34(17): p. 2255–2271.

22. Mangiarini, L., et al., Exon 1 of the HD gene with an expanded CAG repeat is sufficient to cause a progressive neurological phenotype in transgenic mice. Cell, 1996. 87(3): p. 493–506.

23. Barbaro, B.A., et al., Comparative study of naturally occurring huntingtin fragments in Drosophila points to exon 1 as the most pathogenic species in Huntington’s disease. Hum Mol Genet, 2015. 24(4): p. 913–25.

24. Scherzinger, E., et al., Self-assembly of polyglutamine-containing huntingtin fragments into amyloid-like fibrils: Implications for Huntington’s disease pathology. Proceedings of the National Academy of Sciences of the United States of America, 1999. 96(8): p. 4604–4609.

25. Scherzinger, E., et al., Huntingtin-encoded polyglutamine expansions form amyloid-like protein aggregates in vitro and in vivo. Cell, 1997. 90(3): p. 549–558.

26. Hollenbach, B., et al., Aggregation of truncated GST-HD exon 1 fusion proteins containing normal range and expanded glutamine repeats. Philosophical Transactions of the Royal Society of London Series B-Biological Sciences, 1999. 354(1386): p. 991–994.

27. Aiken, C.T., et al., Phosphorylation of threonine 3: implications for Huntingtin aggregation and neurotoxicity. J Biol Chem, 2009. 284(43): p. 29427–36.

28. Steffan, J.S., et al., SUMO modification of Huntingtin and Huntington’s disease pathology. Science, 2004. 304(5667): p. 100–104.

29. Thompson, L.M., et al., IKK phosphorylates Huntingtin and targets it for degradation by the proteasome and lysosome. J Cell Biol, 2009. 187(7): p. 1083–99.

30. Gu, X., et al., Serines 13 and 16 are critical determinants of full-length human mutant huntingtin induced disease pathogenesis in HD mice. Neuron, 2009. 64(6): p. 828–40.

31. Atwal, R.S., et al., Kinase inhibitors modulate huntingtin cell localization and toxicity. Nature Chemical Biology, 2011. 7(7): p. 453–460.

32. Di Pardo, A., et al., Ganglioside GM1 induces phosphorylation of mutant huntingtin and restores normal motor behavior in Huntington disease mice. Proceedings of the National Academy of Sciences of the United States of America, 2012. 109(9): p. 3528–3533.

33. Cariulo, C., et al., Phosphorylation of huntingtin at residue T3 is decreased in Huntington’s disease and modulates mutant huntingtin protein conformation. Proc Natl Acad Sci U S A, 2017. 114(50): p. E10809–E10818.

34. Chaibva, M., et al., Acetylation within the First 17 Residues of Huntingtin Exon 1 Alters Aggregation and Lipid Binding. Biophysical Journal, 2016. 111(2): p. 349–362.

35. Branco-Santos, J., et al., Protein phosphatase 1 regulates huntingtin exon 1 aggregation and toxicity. Human Molecular Genetics, 2017. 26(19): p. 3763–3775.

36. Chiki, A., et al., Mutant Exon1 Huntingtin Aggregation is Regulated by T3 Phosphorylation-Induced Structural Changes and Crosstalk between T3 Phosphorylation and Acetylation at K6. Angew Chem Int Ed Engl, 2017. 56(19): p. 5202–5207.

37. DeGuire, S.M., et al., N-terminal Huntingtin (Htt) phosphorylation is a molecular switch regulating Htt aggregation, helical conformation, internalization, and nuclear targeting. Journal of Biological Chemistry, 2018. 293(48): p. 18540–18558.

38. Daldin, M., et al., Polyglutamine expansion affects huntingtin conformation in multiple Huntington’s disease models. Sci Rep, 2017. 7(1): p. 5070.

39. Warner, J.B., et al., Monomeric Huntingtin Exon 1 Has Similar Overall Structural Features for Wild-Type and Pathological Polyglutamine Lengths. Journal of the American Chemical Society, 2017. 139(41): p. 14456–14469.

40. Cariulo, C., et al., Ultrasensitive quantitative measurement of huntingtin phosphorylation at residue S13. Biochemical and biophysical research communications, 2020. 521(3): p. 549–554.

41. Reif, A., et al., Generation of Native, Untagged Huntingtin Exon1 Monomer and Fibrils Using a SUMO Fusion Strategy. J Vis Exp, 2018(136).

42. Binukumar, B., et al., Profiling of p5, a 24 amino acid inhibitory peptide derived from the CDK5 activator, p35 CDKR1 against 70 protein kinases. Journal of Alzheimer’s Disease, 2016. 54(2): p. 525–533.

43. Hegde, R.N., et al., TBK1 regulates autophagic clearance of soluble mutant huntingtin and inhibits aggregation/toxicity in different models of Huntington’s disease. bioRxiv, 2019: p. 869586.

44. Dan, I., N.M. Watanabe, and A. Kusumi, The Ste20 group kinases as regulators of MAP kinase cascades. Trends in cell biology, 2001. 11(5): p. 220–230.

45. Bustamante, M.B., et al., Detection of huntingtin exon 1 phosphorylation by Phos-Tag SDS-PAGE: Predominant phosphorylation on threonine 3 and regulation by IKKbeta. Biochem Biophys Res Commun, 2015. 463(4): p. 1317–22.

46. Lin, H.K., et al., Fibril polymorphism affects immobilized non-amyloid flanking domains of huntingtin exon1 rather than its polyglutamine core. Nat Commun, 2017. 8: p. 15462.

47. Boatz, J.C., et al., Protofilament Structure and Supramolecular Polymorphism of Aggregated Mutant Huntingtin Exon 1. J Mol Biol, 2020.

48. Caron, N.S., et al., Polyglutamine domain flexibility mediates the proximity between flanking sequences in huntingtin. Proceedings of the National Academy of Sciences of the United States of America, 2013. 110(36): p. 14610–14615.

49. Rockabrand, E., et al., The first 17 amino acids of Huntingtin modulate its sub-cellular localization, aggregation and effects on calcium homeostasis. Hum Mol Genet, 2007. 16(1): p. 61–77.

50. Duim, W.C., et al., Super-resolution fluorescence of huntingtin reveals growth of globular species into short fibers and coexistence of distinct aggregates. ACS chemical biology, 2014. 9(12): p. 2767–2778.

51. Herrera, F., et al., Visualization of cell-to-cell transmission of mutant huntingtin oligomers. PLoS currents, 2011. 3.

52. Wang, H., et al., Effects of overexpression of huntingtin proteins on mitochondrial integrity. Hum Mol Genet, 2009. 18(4): p. 737–52.

53. Cao, F., et al., Nuclear aggregation of huntingtin is not prevented by deletion of chaperone Hsp104. Biochimica et Biophysica Acta (BBA)-Molecular Basis of Disease, 2001. 1537(2): p. 158–166.

54. DiGiovanni, L.F., et al., Huntingtin N17 domain is a reactive oxygen species sensor regulating huntingtin phosphorylation and localization. Hum Mol Genet, 2016.

55. Herrera, F. and J. Branco-Santos, Threonine 3 regulates Serine 13/16 phosphorylation in the huntingtin exon 1. Matters Select, 2019. 5(5): p. e201905000005.

56. Chiki, A., et al., Mutant Exon1 Huntingtin Aggregation is Regulated by T3 Phosphorylation-Induced Structural Changes and Crosstalk between T3 Phosphorylation and Acetylation at K6. Angew Chem Int Ed Engl, 2017.

57. Baias, M., et al., Structure and Dynamics of the Huntingtin Exon-1 N-Terminus: A Solution NMR Perspective. Journal of the American Chemical Society, 2017. 139(3): p. 1168–1176.

58. Newcombe, E.A., et al., Tadpole-like Conformations of Huntingtin Exon 1 Are Characterized by Conformational Heterogeneity that Persists regardless of Polyglutamine Length. J Mol Biol, 2018. 430(10): p. 1442–1458.

59. Urbanek, A., et al., Flanking Regions Determine the Structure of the Poly-Glutamine in Huntingtin through Mechanisms Common among Glutamine-Rich Human Proteins. Structure, 2020.

60. Urbanek, A., et al., A general strategy to access structural information at atomic resolution in polyglutamine homorepeats. Angewandte Chemie International Edition, 2018. 57(14): p. 3598–3601.

61. Eliezer, D., et al., Conformational properties of α-synuclein in its free and lipid-associated states. Journal of molecular biology, 2001. 307(4): p. 1061–1073.

62. Marley, J., M. Lu, and C. Bracken, A method for efficient isotopic labeling of recombinant proteins. Journal of biomolecular NMR, 2001. 20(1): p. 71–75.

63. Hendus-Altenburger, R., et al., Random coil chemical shifts for serine, threonine and tyrosine phosphorylation over a broad pH range. Journal of biomolecular NMR, 2019. 73(12): p. 713–725.

64. Liokatis, S., et al., Simultaneous detection of protein phosphorylation and acetylation by high-resolution NMR spectroscopy. Journal of the American Chemical Society, 2010. 132(42): p. 14704–14705.

65. Kosten, J., et al., Efficient modification of alpha-synuclein serine 129 by protein kinase CK1 requires phosphorylation of tyrosine 125 as a priming event. ACS chemical neuroscience, 2014. 5(12): p. 1203–1208.

66. Chiki, A., et al., Mutant Exon1 Huntingtin Aggregation is Regulated by T3 Phosphorylation-Induced Structural Changes and Crosstalk between T3 Phosphorylation and Acetylation at K6. Angewandte Chemie-International Edition, 2017. 56(19): p. 5202–5207.

67. Froger, A. and J.E. Hall, Transformation of plasmid DNA into E. coli using the heat shock method. J Vis Exp, 2007(6): p. 253.

68. Wishart, D.S. and B.D. Sykes, The 13 C chemical-shift index: a simple method for the identification of protein secondary structure using 13 C chemical-shift data. Journal of biomolecular NMR, 1994. 4(2): p. 171–180.

