## Supporting information for "Site-specific phosphorylation of Huntingtin exon 1 recombinant proteins enabled by the discovery of novel kinases"

and Hilal A. Lashuel *[1]

[1] Laboratory of Molecular and Chemical Biology of Neurodegeneration, School of Life Sciences, Brain Mind Institute, Ecole Polytechnique Fédérale de Lausanne (EPFL), CH-1015 Lausanne, Switzerland.

[2] Protein Production and Structure Core Facility and Laboratory for Biomolecular Modeling, Ecole Polytechnique Fédérale de Lausanne (EPFL) and Swiss Institute of Bioinformatics (SIB), CH-1015 Lausanne, Switzerland.

*To whom correspondence should be addressed: Hilal A. Lashuel, Laboratory of Molecular and Chemical Biology of Neurodegeneration, Brain Mind Institute, Station 19, Ecole Polytechnique Fédérale de Lausanne, CH 1015 Lausanne, Switzerland. Tel: +4121 6939691; Fax: +4121 6931780;

**Table S1**. Expected and observed molecular weights for all the protein generated in this study.

| Proteins | Expected molecular weight (Da) | Observed molecular weight (Da) |
|---|---|---|
| Httex1-23Q | 9943 | 9943 |
| Httex1-43Q | 12505 | 12504; 12509; 12518 ( MALDI); 12525 (MALDI) |
| Httex1-23Q pT3 | 10023 | 10023;10022 |
| Httex1-43Q pT3 | 12585 | 12589;12594 (MALDI); 12604 (MALDI) |
| Httex1-23Q pS13/pS16 | 10103 | 10103;10102.53 |
| Httex1-43Q pS13/pS16 | 12665 | 12665; 12672 (MALDI); 12681 (MALDI) |
| Httex1-23Q 13C/15N | 10516 | 10503.7 |
| Httex1-43Q 13C/15N | 13218 | 13203.4 |
| Httex1-23Q pT3 13C/15N | 10596 | 10583.2 |
| Httex1-43Q pT3 13C/15N | 13298 | 13283.3 |
| Httex1-23Q pS13/pS16 13C/15N | 10676 | 10664.4 |
| Httex1-43Q pS13/pS16 13C/15N | 13378 | 13363.4 |
| Httex1-23Q P90C | 9949 | 9948.07 |
| Httex1-43Q P90C | 12511 | 12511.2 |
| Httex1-23Q pT3 P90C | 10029 | 10029.1 |
| Httex1-43Q pT3 P90C | 12591 | 12591.7 |
| Httex1-23Q pS13/pS16 P90C | 10109 | 10109 |
| Httex1-43Q pS13/pS16 P90C | 12671 | 12671.6 |
| Httex1-23Q pT3 P90C ATTO | 10661 | 10661.9 |
| Httex1-43Q pT3 P90C ATTO | 13223 | 13224.4 |
| Httex1-23Q pS13/pS16 P90C ATTO | 10741 | 10742 |
| Httex1-43Q pS13/pS16 P90C ATTO | 13303 | 13304.9 |

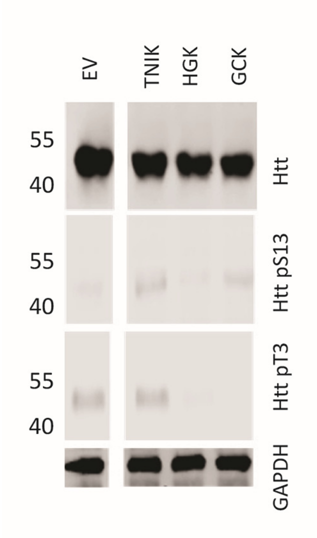

**Figure S1**. Coexpression of Httex1-16Q-eGFP in HEK 293 cells with TNIK, HGK or GCK. Representative Western-blot of Htt, Htt pT3 and Htt pS13 upon co-expression of Httex1-16Q-eGFP with the indicated kinases for 48 hours in HEK 293 cells.

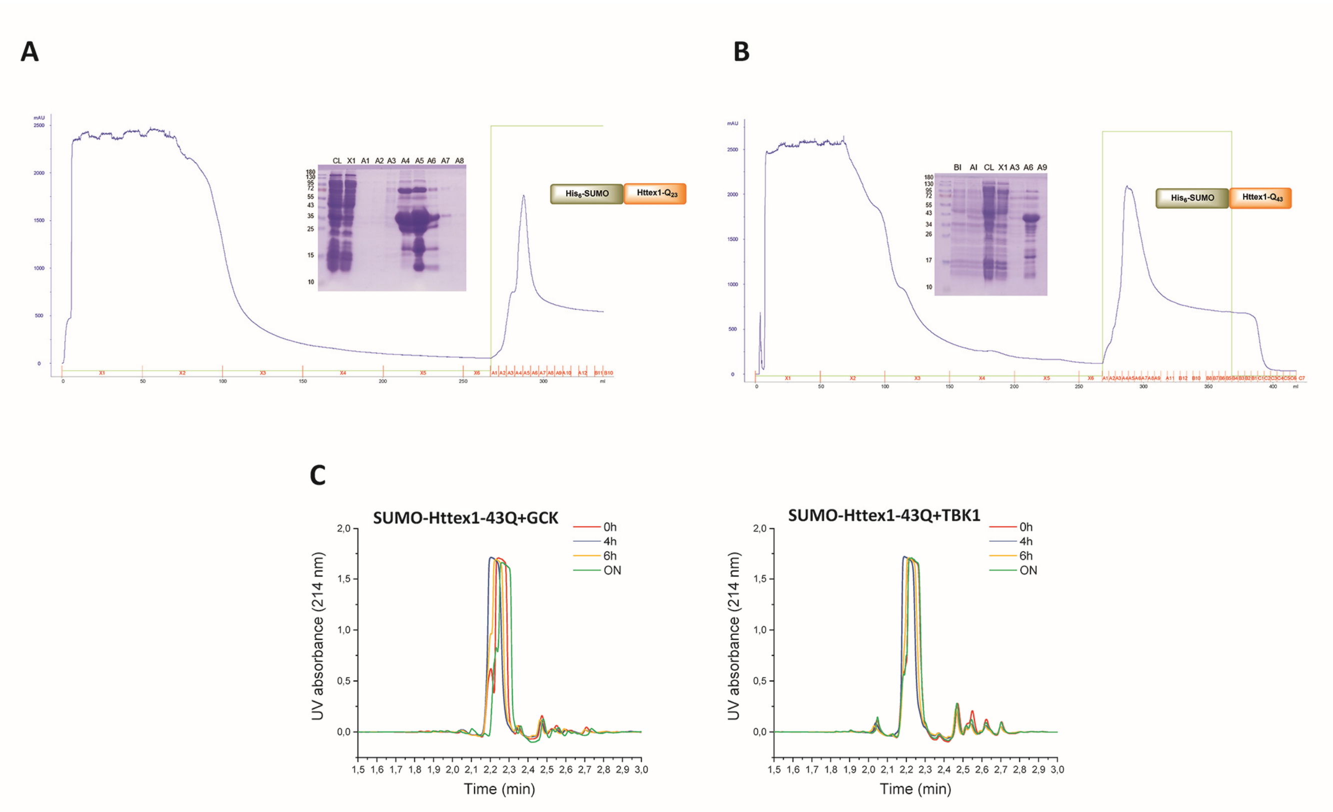

**Figure S2. (A)** Representative chromatogram of the IMAC purification of SUMO-Httex1-23Q, and **(B)** Representative chromatogram of the IMAC purification of SUMO-Httex1-43Q with the analysis of the fractions by SDS-PAGE. **(C)** UPLC spectra of the phosphorylation reaction of SUMO-Httex1-43Q by GCK or TBK1 over-time.

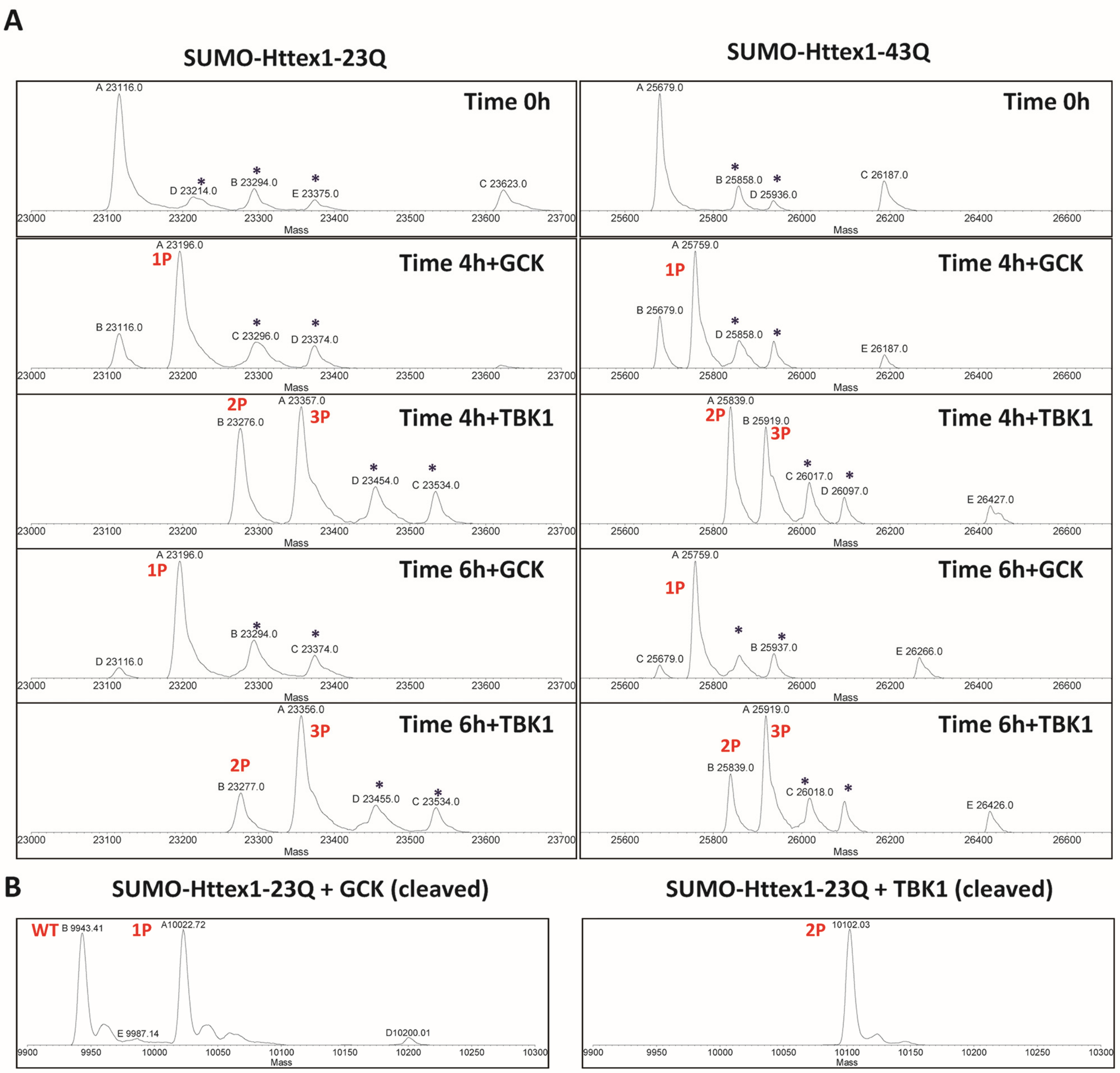

**Figure S3. (A)** phosphorylation of SUMO-Httex1-23Q and SUMO-Httex1-43Q by TBK1 or GCK over-time, followed by ESI/MS (stars are TFA adducts). **(B)** ESI/MS analysis of the phosphorylation reaction of SUMO-Httex1-23Q by GCK (left) and SUMO-Httex1-23Q byTBK1 (right) after SUMO tag cleavage by ULP1**.**

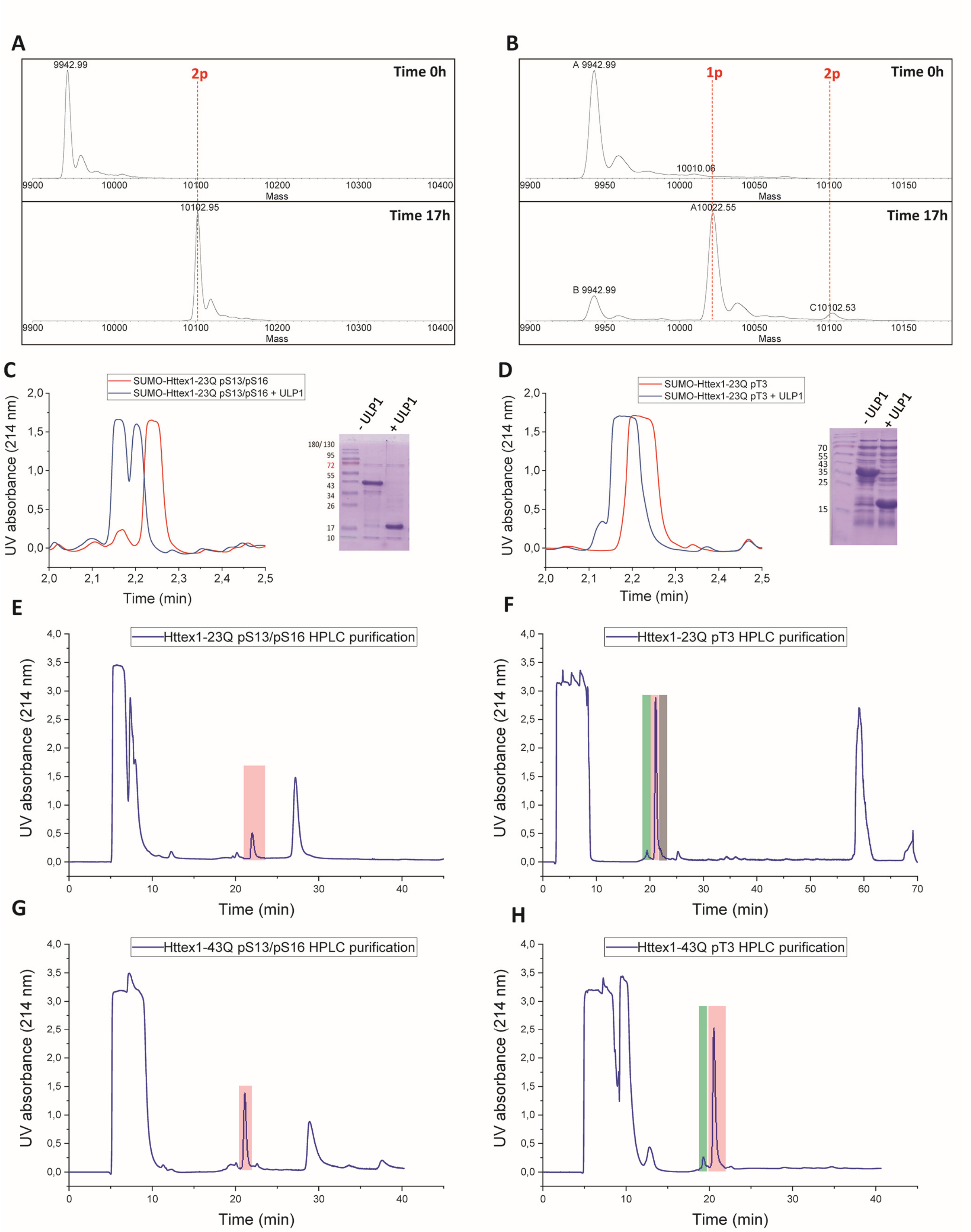

**Figure S4**. Monitoring of the phosphorylation reaction of SUMO-Httex1-23Q with TBK1 **(A)** and GCK **(B)** by ESI/MS after analytical SUMO tag cleavage by ULP1. Monitoring of the cleavage of the SUMO tag by ULP1 from SUMO-Httex1-23Q phosphorylation by TBK1 **(C)** or GCK **(D)**. **(E-H)** RP-HPLC chromatogram for the purification of Httex1-23Q pS13/pS16 **(E)**, Httex1-23Q pT3 **(F)**, Httex1-43Q pS13/pS16 **(G)**, Httex1-43Q pT3 **(H)**, Red: proteins of interest, Green: Httex1 pT3 and pS13, Grey: Unphosphorylated.

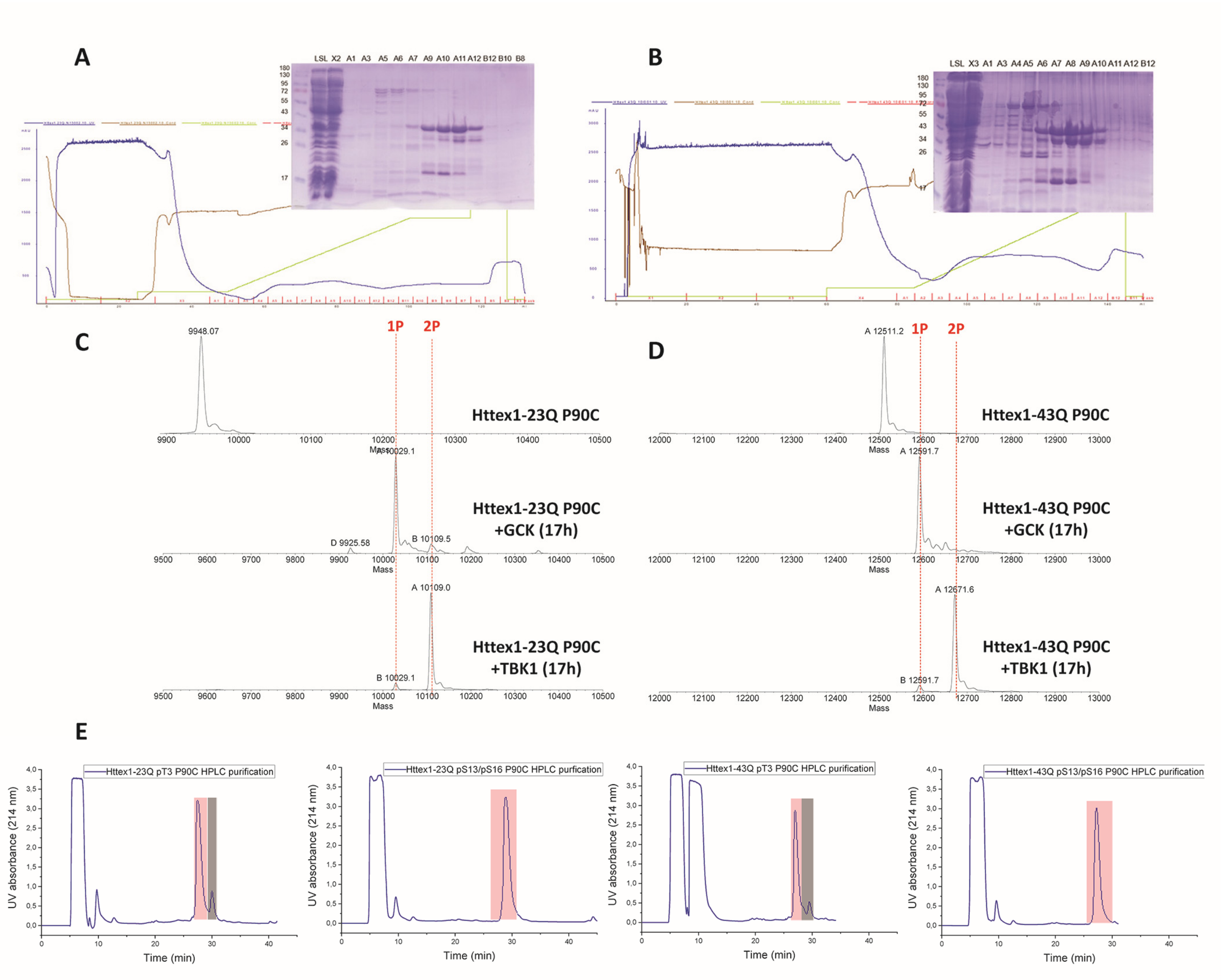

**Figure S5. (A)** Representative chromatogram of the IMAC purification of SUMO-Httex1-23Q P90C, and **(B)** Representative chromatogram of the IMAC purification of SUMO-Httex1-43Q P90C with the analysis of the fractions by SDS-PAGE. **(C)** Monitoring of the phosphorylation reaction by GCK and TBK1 of SUMO-Httex1-23Q P90C by ESI/MS after analytical SUMO tag cleavage by ULP1 **(D)** Monitoring of the phosphorylation reaction by GCK and TBK1 of SUMO-Httex1-43Q P90C by ESI/MS after analytical SUMO tag cleavage by ULP1. **(E)** RP-HPLC purification chromatogram of Httex1-23Q pT3 P90C, Httex1-23Q pS13/pS16 P90C, Httex1-43Q pT3 P90C and Httex1-43Q pS13/pS16 P90C. Red: proteins of interest, Grey: Unphosphorylated.

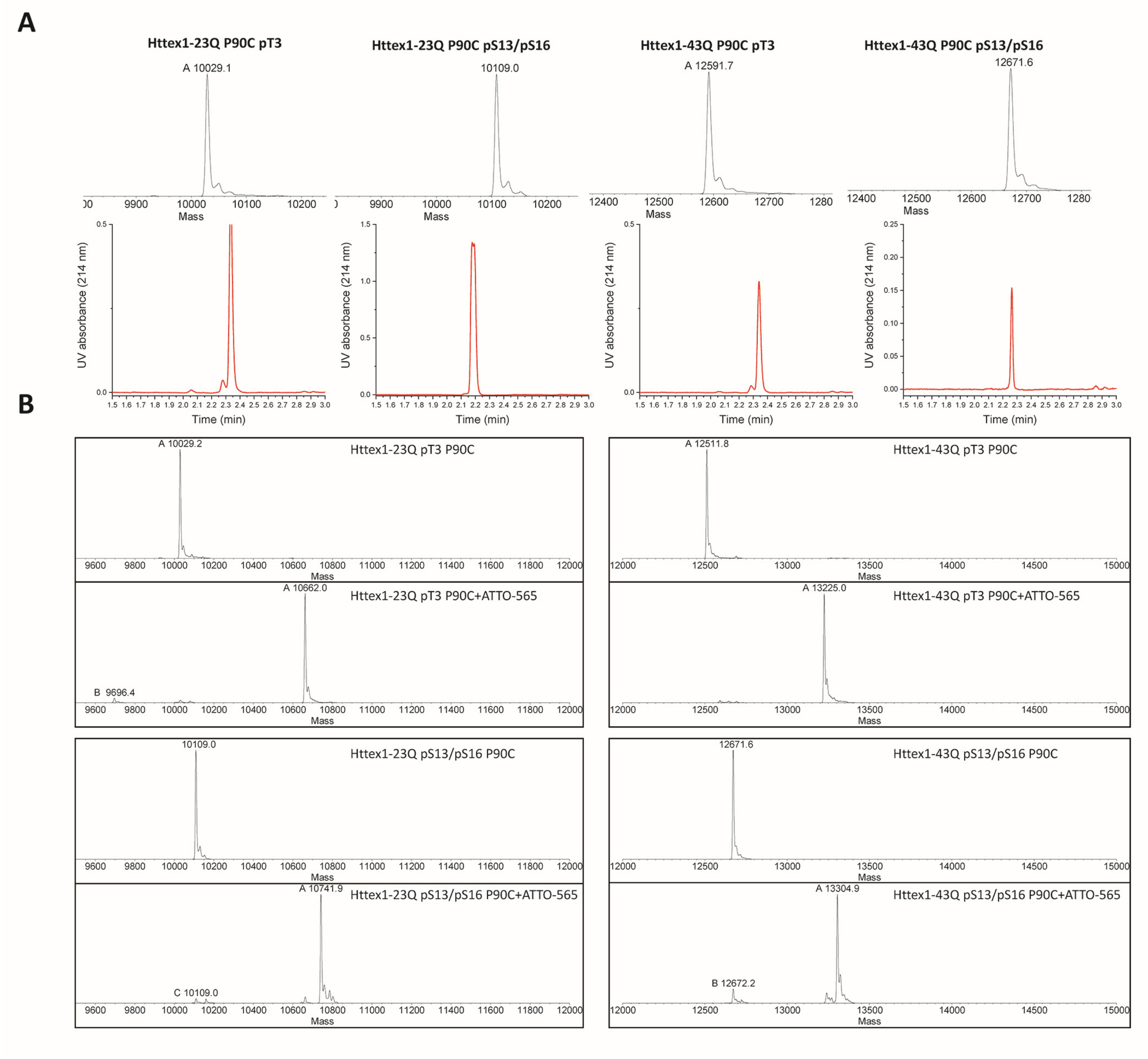

**Figure S6. (A)** Final characterization by ESI/MS ( upper panel) and UPLC (lower panel) of Httex1-23Q pT3 P90C, Httex1-43Q pT3 P90C, Httex1-23Q pS13/pS16 P90C or Httex1-43Q pS13/pS16 P90C and the monitoring of their labelling by ATTO-565-maleimide by ESI/MS **(B)**

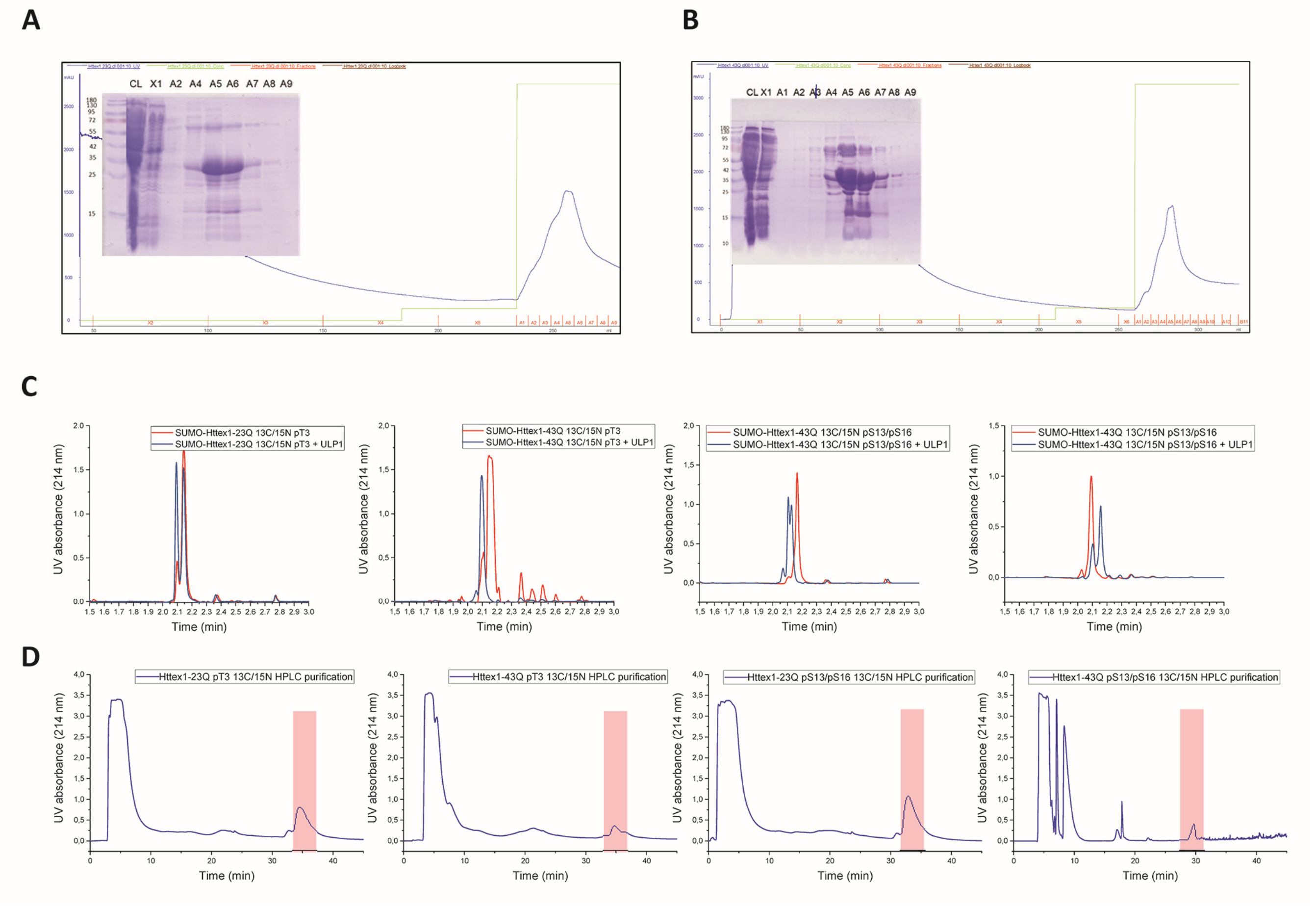

**Figure S7.** Representative chromatogram of the IMAC purification of SUMO-Httex1-23Q $^{13}C/^{15}N$, and **(B)** Representative chromatogram of the IMAC purification of SUMO-Httex1-43Q $^{13}C/^{15}N$ with the analysis of the fractions by SDS-PAGE. **(C)** UPLC Analysis of the cleavage of the SUMO tag by ULP1 from the SUMO-Httex1-Qn $^{13}C/^{15}N$ phosphorylated with GCK or TBK1. **(D)** HPLC purification of Httex1-23Q pT3 $^{13}C/^{15}N$, Httex1-43Q pT3 $^{13}C/^{15}N$, Httex1-23Q pS13/pS16 $^{13}C/^{15}N$ or Httex1-43Q pS13/pS16 $^{13}C/^{15}N$. Red: proteins of interest.

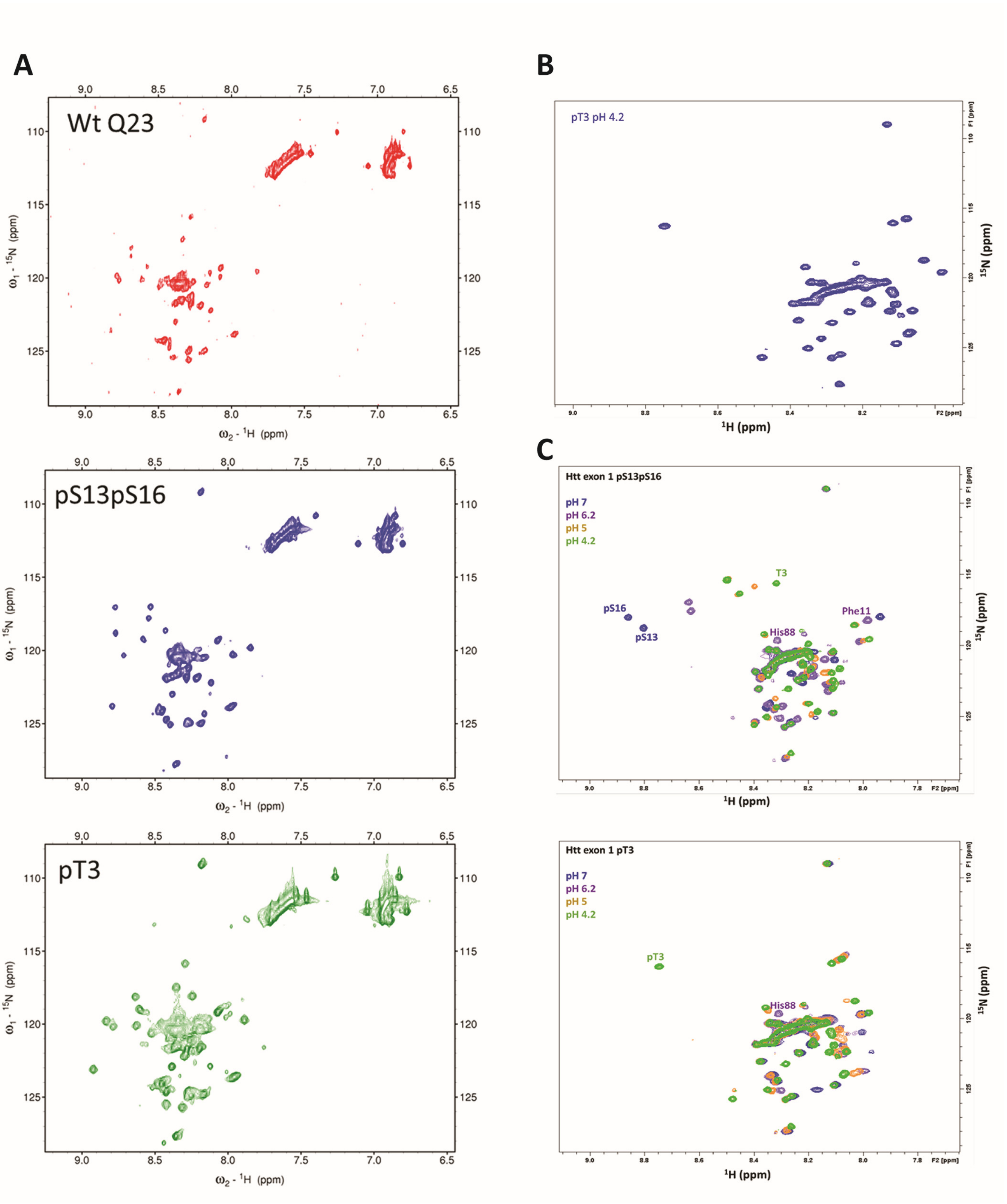

**Figure S8**. **(A)** $^{1}$H,$^{15}$N HSQC spectra of unmodified, pS13pS16 and pT3 Httex1-23Q proteins in 20% trifluoroethanol, showing an increased chemical shift dispersion that likely reflects helical structuring occurring in all three species. **(B)** $^{1}$H,$^{15}$N HSQC spectrum of Httex1-23Q pT3 acquired at pH 4.2 and 25°C. **(C)** $^{1}$H,$^{15}$N HSQC spectra of Httex1-23Q pT3 and Httex1 pS13/pS16 proteins at pH 7, 6.2, 5 and 4.2, with the assignments of residues discussed in the text.

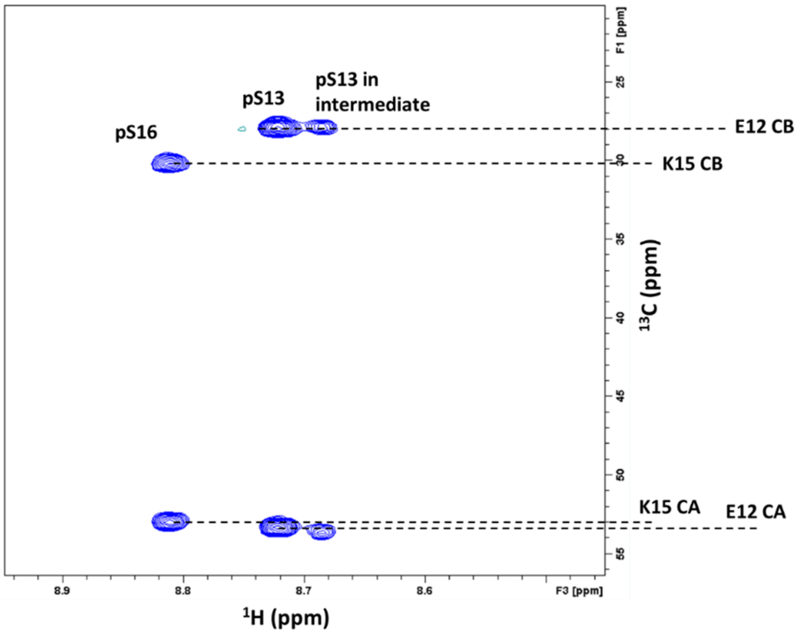

**Figure S9**. $^{1}$H,$^{13}$C plane of a CBCA(CO)NH spectrum collected after 21 hours of reaction between Httex1-23Q and TBK1. The $^{15}$N frequency was not evolved, allowing an experimental acquisition time of 20 minutes to minimize changes occurring while the spectrum was acquired. The $^{13}$C chemical shifts of the residues preceding Ser13 and Ser16 are shown to support that the phosphorylation intermediate corresponds to Ser13, especially given the clear distinction between Glu12 and Lys15 CB shifts.

**Table S2.** H,N,CA,CB assignments for human Htt ex1 unmodified and pS13pS16 in our working conditions, pH 7, 25 ºC

| | | **Unmodified Htt exon 1** | | | | | | **pS13pS16 Htt exon 1** | | | |
|---|---|---|---|---|---|---|---|---|---|---|---|
| **AA** | **No.** | **H** | **N** | **CA** | **CB** | **AA** | **No.** | **H** | **N** | **CA** | **CB** |
| Thr | 3 | | | 59.16 | 67.02 | Thr | 3 | | | 59.16 | 67.02 |
| Leu | 4 | 8.41 | 124.6 | 52.82 | 39.3 | Leu | 4 | 8.38 | 125.2 | 52.82 | 39.3 |
| Glu | 5 | 8.36 | 121.7 | 54.56 | 27.36 | Glu | 5 | 8.34 | 122.1 | 54.04 | 27.36 |
| Lys | 6 | | | 54.06 | 29.89 | Lys | 6 | 8.2 | 122.1 | 53.59 | 30.07 |
| Leu | 7 | 8.06 | 122.3 | 52.76 | 39.38 | Leu | 7 | 8.12 | 123.1 | 52.43 | 39.38 |
| Met | 8 | | | 52.63 | 29.85 | Met | 8 | 8.22 | 121.2 | 52.44 | 29.86 |
| Lys | 9 | 8.09 | 121.7 | 54.02 | 30.36 | Lys | 9 | 8.12 | 122.7 | 53.4 | 30.36 |
| Ala | 10 | 8.07 | 123.9 | 50.34 | 16.34 | Ala | 10 | 8.24 | 125.1 | 49.91 | 16.34 |
| Phe | 11 | 8.03 | 118.6 | 55.8 | 36.85 | Phe | 11 | 7.94 | 118 | 54.72 | 36.85 |
| Glu | 12 | 8.12 | 121.4 | 54.49 | 27.28 | Glu | 12 | 8.12 | 122.1 | 53.4 | 27.92 |
| Ser | 13 | 8.08 | 115.8 | 56.21 | 60.84 | pSer | 13 | 8.73 | 118.6 | 54.99 | 63.02 |
| Leu | 14 | 8.03 | 123.5 | 53.22 | 39.35 | Leu | 14 | 8.35 | 124.1 | 52.57 | 39.35 |
| Lys | 15 | 8.06 | 120.6 | 54.39 | 30.11 | Lys | 15 | 8.22 | 122.5 | 53.13 | 30.11 |
| Ser | 16 | 8.07 | 115.5 | 56.75 | 60.76 | pSer | 16 | 8.83 | 117.9 | 55.79 | 62.52 |
| Phe | 17 | 8.12 | 122.1 | 56.52 | 36.37 | Phe | 17 | 8.24 | 121.7 | 56.52 | 36.37 |
| Gln | 18 | 8.2 | 120 | 53.94 | 26.13 | Gln | 18 | 8.05 | 120.8 | 53.94 | 26.13 |
| Gln | 19 | 8.11 | 119.9 | 54.3 | 25.93 | Gln | 19 | 8.07 | 120.3 | 54.3 | 25.93 |
| Gln | 38 | | | 53.23 | 26.62 | Gln | 38 | | | 53.23 | 26.62 |
| Gln | 39 | 8.36 | 121.4 | 52.98 | 26.83 | Gln | 39 | 8.36 | 121.4 | 52.98 | 26.83 |
| Gln | 40 | 8.37 | 122.8 | 50.84 | 26.07 | Gln | 40 | 8.37 | 122.9 | 50.84 | 26.07 |
| | | | | 60.16 | 29.18 | | | | | | |
| Gln | 52 | 8.32 | 120.2 | 52.63 | 26.89 | Gln | 52 | | | 52.68 | 26.87 |
| Leu | 53 | 8.28 | 125.3 | 50.21 | 38.69 | Leu | 53 | 8.28 | 125.3 | 50.21 | 38.69 |
| Pro | 58 | | | 60.25 | 29.14 | Pro | 58 | | | 60.25 | 29.14 |
| Gln | 59 | 8.34 | 120.3 | 53.09 | 26.75 | Gln | 59 | 8.34 | 120.3 | 53.09 | 26.75 |
| Ala | 60 | 8.29 | 125.6 | 49.57 | 16.36 | Ala | 60 | 8.29 | 125.6 | 49.57 | 16.36 |
| Pro | 62 | | | 60.39 | 29.27 | Pro | 62 | | | 60.39 | 29.27 |
| Leu | 63 | 8.23 | 122.2 | 52.18 | 39.44 | Leu | 63 | 8.23 | 122.2 | 52.18 | 39.44 |
| Leu | 64 | 8.11 | 124.5 | 49.85 | 38.68 | Leu | 64 | 8.11 | 124.6 | 49.85 | 38.68 |
| Pro | 65 | | | 60.12 | 29.3 | Pro | 65 | | | 60.12 | 29.3 |
| Gln | 66 | 8.4 | 121.6 | 50.53 | 26.11 | Gln | 66 | 8.4 | 121.6 | 50.53 | 26.11 |
| Pro | 78 | | | 60.23 | 29.26 | Pro | 78 | | | 60.42 | 29.26 |
| Gly | 79 | 8.13 | 108.7 | 41.55 | | Gly | 79 | 8.13 | 108.8 | 41.55 | |
| Pro | 80 | | | 60.01 | 29.28 | Pro | 80 | | | 60.39 | 29.28 |
| Ala | 81 | 8.31 | 124.2 | 49.54 | 16.37 | Ala | 81 | 8.31 | 124.2 | 49.54 | 16.37 |
| Val | 82 | 8.02 | 119.7 | 59.27 | 30.01 | Val | 82 | 8.02 | 119.7 | 59.27 | 30.01 |
| Ala | 83 | 8.29 | 127.8 | 49.47 | 16.34 | Ala | 83 | 8.28 | 127.9 | 49.47 | 16.34 |
| Glu | 84 | | | 53.36 | 27.7 | Glu | 84 | 8.34 | 123.9 | 53.36 | 27.7 |
| Glu | 85 | 8.34 | 123.9 | 51.49 | 26.96 | Glu | 85 | | | 51.49 | 26.96 |
| Pro | 86 | | | 60.37 | 29.18 | Pro | 86 | | | 60.37 | 29.18 |

| | | | | | | | | | | | |
|---|---|---|---|---|---|---|---|---|---|---|---|
| Leu | 87 | 8.19 | 121.6 | 52.47 | 39.58 | Leu | 87 | 8.18 | 121.5 | 52.47 | 39.58 |
| His | 88 | 8.15 | 120.2 | 53.06 | 27.86 | His | 88 | 8.22 | 119.9 | 52.41 | 27.39 |
| Arg | 89 | 8.13 | 124.9 | 50.91 | 27.3 | Arg | 89 | 8.2 | 124.9 | 50.91 | 27.64 |
